# Mechanically induced localization of SECONDARY WALL INTERACTING bZIP is associated with thigmomorphogenic and secondary cell wall gene expression

**DOI:** 10.1101/2021.02.03.429573

**Authors:** Joshua H. Coomey, Kirk J.-M. MacKinnon, Ian W. McCahill, Bahman Khahani, Pubudu P. Handakumbura, Gina M. Trabucco, Jessica Mazzola, Nicole A. Leblanc, Rithany Kheam, Miriam Hernandez-Romero, Kerrie Barry, Lifeng Liu, Ji E. Lee, John P. Vogel, Ronan C. O’Malley, James J. Chambers, Samuel P. Hazen

## Abstract

Plant growth requires the integration of internal and external cues, perceived and transduced into a developmental program of cell division, elongation, and wall thickening. Mechanical forces contribute to this regulation, and thigmomorphogenesis typically includes reducing stem height, increasing stem diameter, and a canonical transcriptomic response. We present data on a bZIP transcription factor involved in this process in grasses. *Brachypodium distachyon* SECONDARY WALL INTERACTING bZIP (SWIZ) protein translocated into the nucleus following mechanostimulation. Classical touch responsive genes were upregulated in *B. distachyon* roots following touch, including significant induction of the glycoside hydrolase 17 family, which may be unique to grass thigmomorphogenesis. SWIZ protein binding to an E-box variant in exons and introns was associated with immediate activation followed by repression of gene expression. *SWIZ* overexpression resulted in plants with reduced stem and root elongation. These data further define plant touch-responsive transcriptomics and physiology, offering insights to grass mechanotranduction dynamics.

## INTRODUCTION

Forces both internal and external to a cell influence growth. Turgor pressure in conjunction with anisotropic cell wall dynamics direct plant cell shape and expansion. Force perception between neighboring cells is critical in the development and maintenance of tissue form and function, such as the interlocking pavement cells on the leaf epidermis, or the developmental hotspots in the apical meristem (Hamant et al., 2008; Uyttewaal et al., 2012; Bidhendi et al., 2019). Specific inter-cell forces result in dynamic remodeling of the cortical cytoskeleton, with subsequent changes in cellulose microfibril alignment and alterations to other cell wall components such as pectin methyl esterification (Hamant et al., 2008; Uyttewaal et al., 2012; Bidhendi and Geitmann, 2018; Altartouri et al., 2019; Bidhendi et al., 2019). The classic hallmarks of touch responsive growth, or thigmomorphogenesis, include reduced plant height, increased radial growth in plants with a cambial meristem, increased branching, and delayed flowering time (Jaffe, 1973; Biro et al., 1980; Jaffe et al., 1980; Braam, 2004). These attributes have been leveraged by farmers for hundreds of years. As early as 1680, records show Japanese farmers tread on young wheat and barley seedlings to elicit increased branching, spikes per plant, and grain weight per plant, along with stronger roots (Iida, 2014). This practice, known as mugifumi, continues today with mechanized rollers. Thigmomorphogenesis in belowground tissues have also been studied to some extent, with the impact of stiffer substrates eliciting changes in root length and straightness, with the implication of hormonal signaling pathways in mediating this response (Lourenço et al., 2015; Lee et al., 2019; Nam et al., 2020).

Mechanical stimulus can significantly remodel gene expression (Braam and Davis, 1990; Braam, 2004; Lee et al., 2005). The so-called *TOUCH* (*TCH*) genes in *Arabidopsis thaliana*, encode calmodulin (*AtTCH1*/*AtCaM2*), calmodulin-like proteins (*AtTCH2*/*AtCML24*, *AtTCH3*/*CML12*), and a xyloglucan endotransglucosylase/hydrolase (*AtTCH4*/*AtXTH22*) (Braam and Davis, 1990). Touch responsive gene expression patterns often overlap with other stimuli such as dark, cold, and hormone treatment (Polisensky and Braam, 1996; Lee et al., 2005). In addition to calcium binding and signaling, genes related to cell wall modification and a variety of transcription factors and kinases are regulated by mechanical stimulus, as well as genes involved in hormone homeostasis and signaling.

Group I bZIPs are also implicated in mechanosensing. VIRE2-INTERACTING PROTEIN 1 (AtVIP1) and related Group I bZIP proteins translocate from the cytoplasm to the nucleus in response to a variety of biotic and abiotic stimuli, including hypo-osmotic conditions (Tsugama et al., 2012; Tsugama et al., 2014; Tsugama et al., 2016). The Group I bZIP *Nt REPRESSOR OF SHOOT GROWTH* (*NtRSG*) in tobacco plays a role in maintaining GA homeostasis, wherein it translocates to the nucleus in response to cellular bioactive GA levels (Igarashi et al., 2001; Ishida et al., 2008; Fukazawa et al., 2010; Ito et al., 2017). Translocation appears to be dependent on protein phosphorylation, either from MITOGEN ACTIVATED PROTEIN KINASE 3 during pathogen invasion, or via calcium dependent protein kinases. When phosphorylated, Group I bZIPs associate with 14-3-3 proteins in the cytoplasm until phosphatase activity releases them for nuclear translocation (Ishida et al., 2004; Ito et al., 2014; Ito et al., 2017; Tsugama et al., 2018; Van Leene et al., 2016).

Secondary cell walls deposited between the plasma membrane and primary cell wall provide mechanical strength in vascular and structural tissues. Secondary walls are made of crystalline cellulose, hemicelluloses, and phenolic lignin polymers. Although functionally similar, secondary walls in monocotyledonous plants have key differences from eudicots, including distinct hemicellulose chemistry and differences in lignin biosynthesis. Grasses also produce mixed-linkage glucans (MLGs), a wall polysaccharide that is rarely found outside the commelinid monocots (Coomey et al., 2020). A tightly controlled network of feed-forward loops regulates the transcription of wall synthesizing enzymes, with NAC family transcription factors activating wall synthesis genes as well as MYB family and other transcription factors that further promote secondary wall synthesis (McCahill and Hazen, 2019). These networks are similar between grasses and eudicots, with some components in each that have yet to be described in the other (Coomey et al., 2020). There are currently no bZIP family members in any secondary wall regulatory model.

Grasses employ a fundamentally different growth mechanism than eudicots, as they lack a cambium layer, and thus no lateral meristem. Stem elongation comes from division and elongation in a discrete series of intercalary meristems, called nodes, with one internode region elongating and pushing up subsequent nodes. Detailed studies of thigmomorphogenesis have been conducted almost exclusively in eudicots. However, recent work in the model cereal grass *Brachypodium distachyon* shows general overlap with conventional eudicot thigmomorphogenesis but with no change in stem diameter and increased time to flower (Gladala-Kostarz et al., 2020).

Thigmomorphogenesis is a widely observed phenomenon that results in reduced height, increased radial growth, and increased branching. The mechanisms behind this form of growth are not yet fully understood, but involve aspects of hormone regulation, Ca^2+^ signaling, Group I bZIP intracellular translocation, and changes in gene expression. Here we describe the transcriptional response to mechanical stimulation and the function of a *B. distachyon* bZIP transcription factor, SECONDARY WALL ASSOCIATED bZIP (Bradi1g17700) and its role in touch response and cell wall biosynthesis.

## RESULTS

### SWIZ is a Group I bZIP transcription factor and candidate cell wall regulator

To identify genes involved in the regulation of secondary cell wall thickening, Trabucco et al., (2013) measured transcript abundance in *B. distachyon* leaf, root, and stem tissue. A gene annotated as a bZIP transcription factor, Bradi1g17700, was highly expressed in root and stem relative to leaf (**Supplemental Figure S1A**). Bradi1g17700 is also a member of a 112 gene coexpression network (**Supplemental Table S1**) that includes genes highly expressed in the peduncle (Sibout et al., 2017). Phylogenetic analysis of Bradi1g17700, hereinafter referred to as SECONDARY WALL INTERACTING bZIP (SWIZ), amino acid sequence shows it to be an ortholog of the *A. thaliana* Group I bZIPs (Jakoby et al., 2002; Dröge-Laser et al., 2018), and closely related to *AtbZIP18* and *AtbZIP52* (**Supplemental Figure S1B, Supplemental File S1**).

### SWIZ translocates into the nucleus in response to mechanical stimulus

Group I bZIPs in *A. thaliana* translocate between the cytosol and nucleus in response to external cellular force. As an ortholog of these proteins, we hypothesized that SWIZ protein may similarly translocate within the cell in response to mechanical force. To test this, the roots of plants overexpressing either *SWIZ* fused to *GFP* (*SWIZ:GFP-OE*) or *GFP* alone (*GFP-OE*) were observed following a mechanical stimulus (**Fig 1A, Supplemental Files 2 and 3**). In control plants, GFP signal was both cytosolic and nuclear, which remained constant over the imaging period (**Fig 1B**). GFP signal was mostly observed in the cytosol in *SWIZ:GFP-OE* plants, but following mechanical stimulus, nuclear GFP signal increased substantially, peaking around 30 min post-stimulus and returned to near basal levels by ∼60 min, while untouched plants showed no change in signal localization (**Fig 1B**).

**Figure 1.**
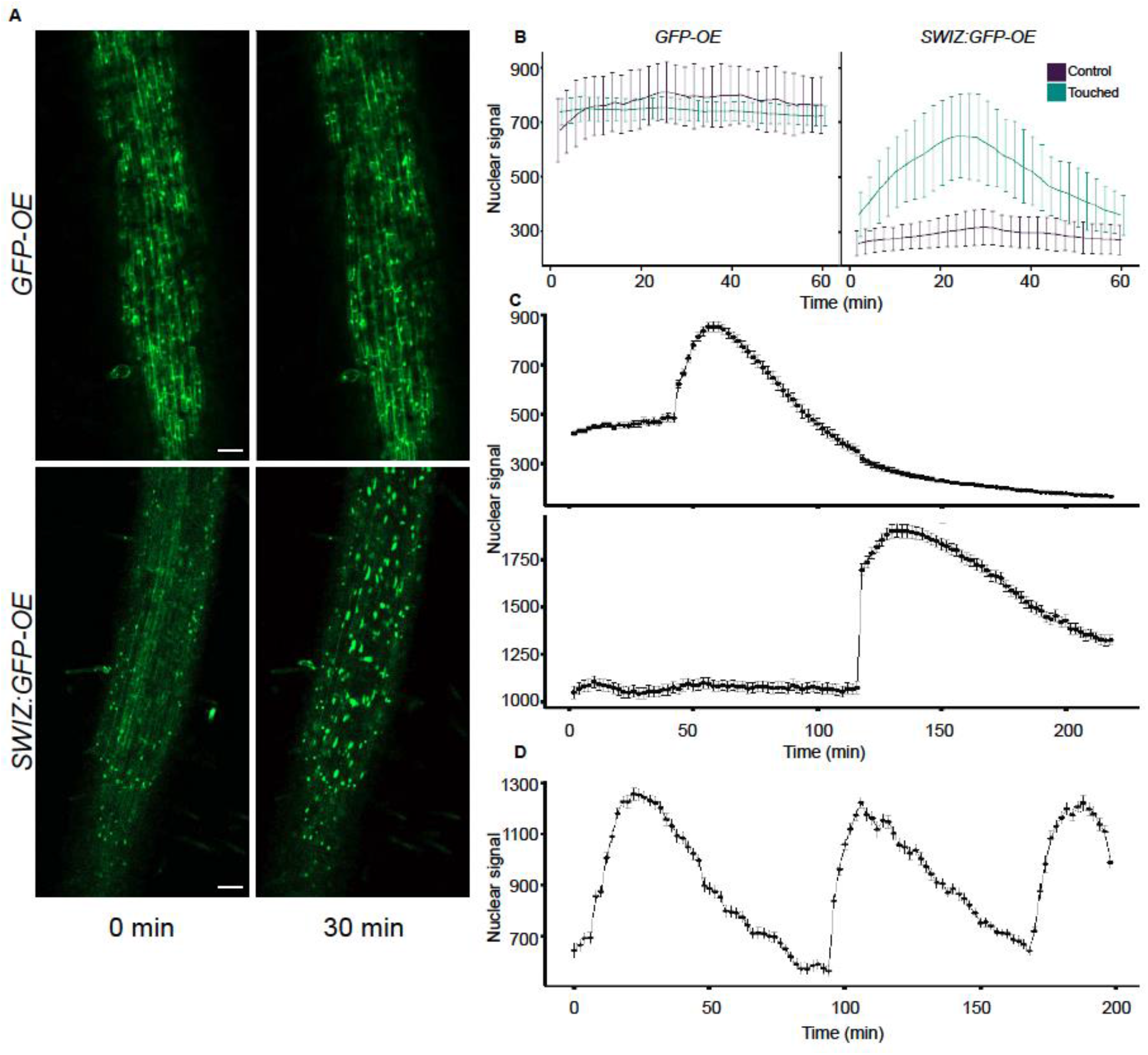
SWIZ translocates to the nucleus in response to mechanical stimulus, specifically in regions directly stimulated. (A) Image of *SWIZ:GFP-OE* and *GFP-OE* roots prior to stimulus and 30 min post stimulus. Roots were observed immediately following mechanical perturbation. (B) Quantification of nuclear signal in control (purple) and touched (teal) conditions for *GFP-OE* (left) and *SWIZ:GFP-OE* (right). n = 14-20 nuclei. (C) SWIZ translocation occurred in the local area of the stimulus. At 30 min, stimulus was applied to an upper region of the root, while at 120 min it was applied to a lower region approximately 3 cm below. n = 109, 184 nuclei respectively for upper and lower regions. Scale bar = 100 µm, (D) *SWIZ:GFP-OE* roots were imaged by confocal microscopy with stimulus applied in the field of view at 0, 90, and 180 min. n = 126 nuclei. (B-D) Images were taken every 2 min. Nuclear GFP signal was quantified in selected nuclei at each time point. The average nuclear GFP signal is represented by the line with error bars indicating standard error of the mean. Scale bar = 100 µm. n = 4-6 plants per treatment.

Touch response to stimulus can saturate at a certain number of treatments (Martin et al., 2010; Leblanc-Fournier et al., 2014; Moulia et al., 2015). To test if SWIZ translocation dynamics varied after repeated treatments, we applied mechanical force to *SWIZ:GFP-OE* roots at regular intervals. A second stimulus was given 90 min after the first, and a third at 180 min. Following each mechanical stimulation, SWIZ consistently translocated from cytoplasm to nucleus (**Fig 1D**). This suggests that SWIZ translocation dynamics are not impacted by repeated stimulus events 90 min apart.

To determine if the signal triggering SWIZ translocation is spread beyond the specifically stimulated region, two regions of the same *SWIZ:GFP-OE* root separated by 3 cm were simultaneously observed. The stimulated region showed typical SWIZ:GFP nuclear signal accumulation and in the region below no translocation was observed (**Fig 1C**). At 120 min, the treatments were reversed, with the lower root region receiving a stimulus while the upper region was unperturbed. The lower region showed SWIZ:GFP nuclear translocation while the upper region did not. Thus, the touch stimulus is localized directly to the perturbed region.

### Transcriptional response to touch

Having established the nuclear translocation of SWIZ in response to touch, we then investigated what effect touch and *SWIZ* overexpression during touch may have on gene expression. We measured transcript abundance by sequencing mRNA from wildtype and *SWIZ-OE* root tissue just prior to touch (0 min) and at 10, 30, and 60 min following touch treatment along the entire length of the root (**Fig 2A, Supplemental Fig S2**). Principal component analysis of transcript abundance shows the greatest amount of variance is attributed to the first principal component, 60%, where samples cluster based on genotype (**Fig 2B**). The second principal component, which accounts for 17% of the variance, distinguished between pre-touch and 10 min following touch, with the last two time points clustering similarly.

**Figure 2.**
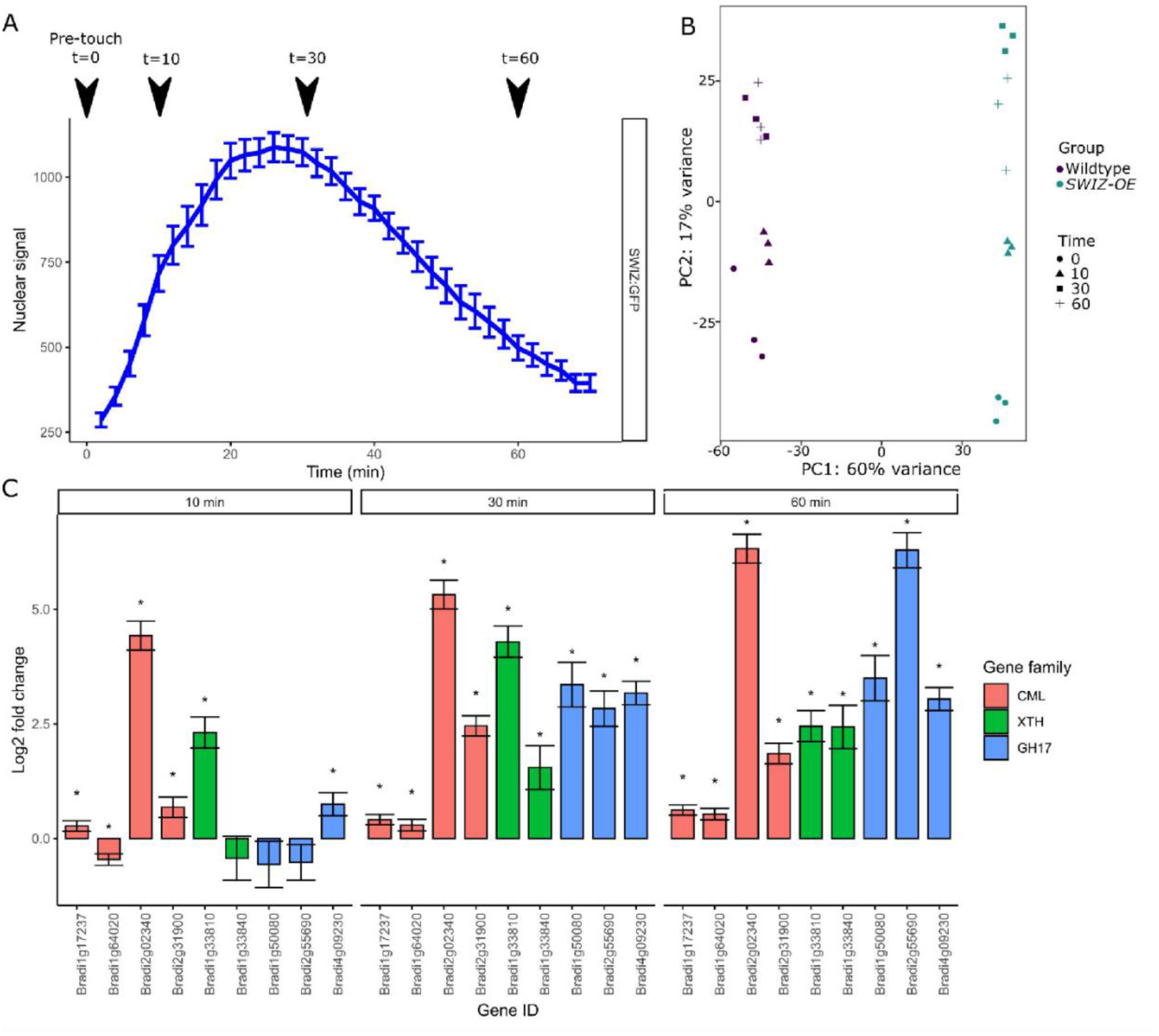
Transcriptome analysis of touch response in *Brachypodium distachyon* roots. (A) Root tissue was sampled just prior to touch (t = 0), and at 10, 30, and 60 min following touch treatment in wildtype and *SWIZ-OE.* (B) Principal component analysis of gene expression across samples shows the greatest difference corresponding to genotype and the second greatest corresponding to time after touch. (C) Canonical, as well as novel touch responsive genes are upregulated in *B. distachyon* following touch. Closest orthologs of the *Arabidopsis thaliana TOUCH* genes encoding calmodulin-like (CML) and xyloglucan endo- transglycosylase/hydrolases (XTH) are upregulated following touch, as are previously unreported members of the glycosyl hydrolase 17 (GH17) family. Full list of gene expression for *TOUCH* and glycoside hydrolases defined by Tyler et al. (2010) in Supplemental Tables S10-12. Significance denoted by * reflecting *q* < 0.1 compared to expression at t = 0, with *q*-values representing Wald test *p*-values adjusted for false discovery rate.

The wildtype transcriptome was massively remodeled in response to touch, with 8,902 transcripts differentially expressed (*q* < 0.1) at 10 min post touch, 5,682 transcripts at 30 min, and 7,672 transcripts at 60 min (**Supplemental Tables S2-4**). Canonical touch-responsive genes (TCH) include calmodulins (CAM), calmodulin-like (CmL), and xyloglucan endo- transglycosylase/hydrolases (XTH) (Braam and Davis, 1990; Lee et al., 2005). Based on homology with *TCH1*, *TCH2*, and *TCH3*, 79 CAM and CmL genes were identified, the majority of which showed upregulation in wildtype plants following touch treatment (**Fig 2C**, **Supplemental Table S11**). Similarly, homology with *TCH4* identified 37 *B. distachyon XTH* and *XTH-like* genes. Of these, 8 genes were upregulated by touch in wildtype plants, including the two with the highest similarity to *TCH4* (Bradi1g33810 and Bradi33840) (**Fig 2C**, **Supplemental Table S12**). We also investigated the effect of touch on other glycosyl hydrolase families (Tyler et al., 2010). GH17 genes were differentially regulated following touch. The GH17 family is broadly described as modifying β-1,3-glucans. Thirty-five of the 53 annotated GH17 members were measured in our data, with 8, 11, and 16 members differentially expressed at 10, 30 and 60 min respectively. Fisher’s exact test determined significant enrichment of GH17 expression at time 60 (*p* = 9.307e-3) (**Fig 2C**, **Supplemental Table S12**). An interactive platform for exploring this *B. distachyon* touched-root transcriptome is available online at (https://hazenlab.shinyapps.io/swiztc/).

### Cell wall genes are downregulated immediately following touch, then induced concurrent with SWIZ nuclear translocation and more strongly in *SWIZ-OE*

Genes related to secondary cell wall synthesis were immediately downregulated in wildtype plants following touch, but were subsequently upregulated (**Fig 3, Supplemental Table S5-10**). Xylan, cellulose, MLG, and callose synthesis associated genes were upregulated 30 and 60 min following touch, while lignin biosynthesis and BAHD acyltransferase genes were upregulated at 60 min. Prior to touch, nearly all cell wall associated genes investigated were significantly downregulated relative to wildtype, then significantly upregulated following touch (**Fig 3**). In *SWIZ-OE*, cell wall-related terms were enriched in paths that had stronger upregulation earlier than the same terms in wildtype (**Supplemental Fig S3A-B**). Network analysis of gene expression patterns over time using the iDREM software showed that in *SWIZ-OE* plants, cell wall genes were more rapidly induced. Notably, cell wall polysaccharide biosynthesis, particularly glucan biosynthesis, were significantly enriched in *SWIZ-OE* path D, while not significantly enriched in wildtype (**Supplemental Table S14**). This is mirrored when looking at specific gene expression, with primary wall CESAs, MLG, and callose synthesis genes all significantly upregulated following touch in *SWIZ-OE*. Thus, cell wall genes were immediately repressed by touch, followed by significant activation, which occurs more rapidly in *SWIZ-OE* plants and is associated with the timing of SWIZ nuclear translocation.

**Figure 3.**
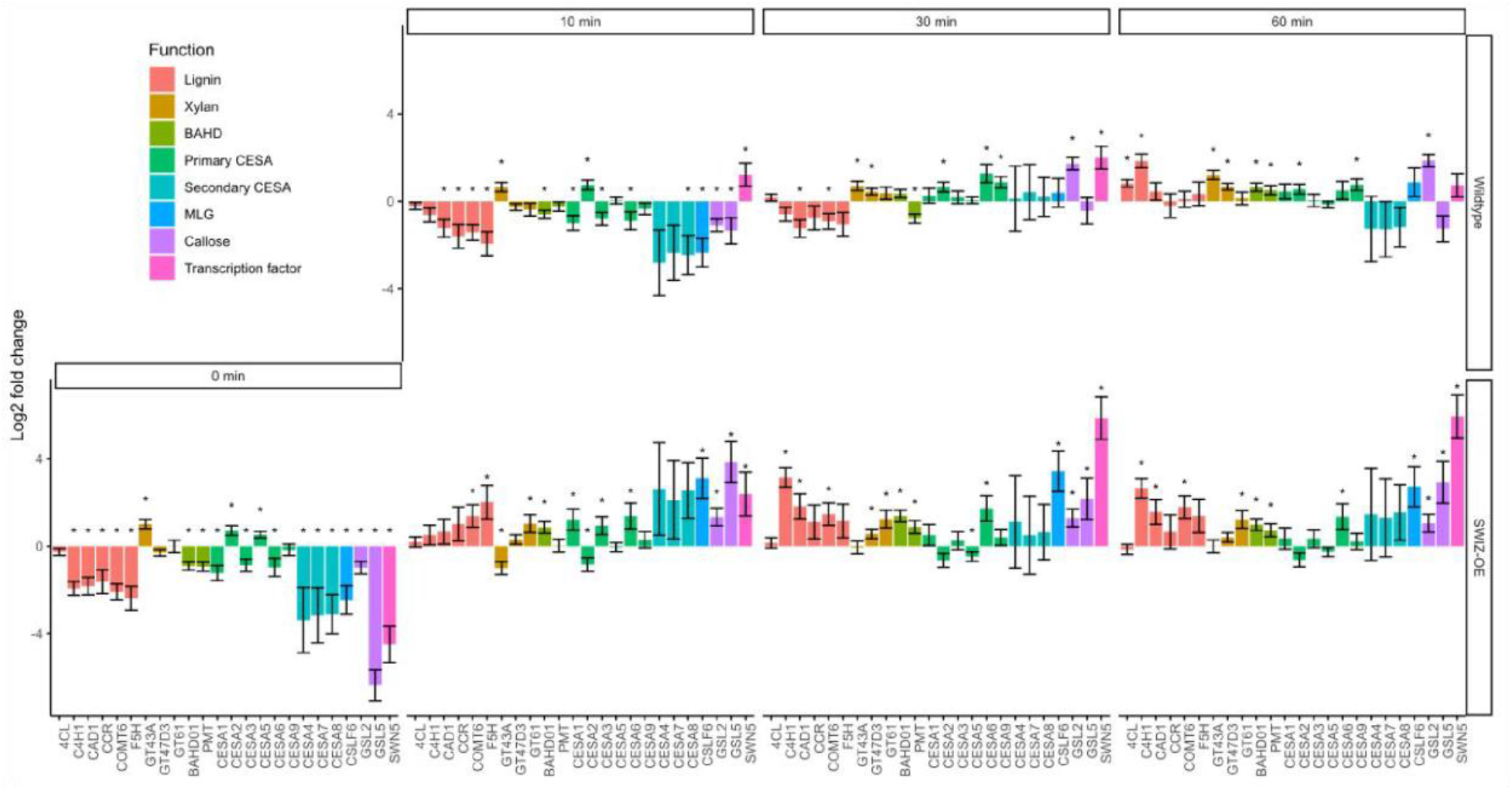
Gene expression analysis of cell wall related genes. Log fold-change of gene expression measured by RNA-seq in wildtype and *SWIZ-OE*, presented as relative to wildtype expression at time 0, pre-touch. Bar color indicates class of cell wall gene. Error bars indicate standard deviation of three biological replicates. Significance denoted by * reflecting *q* < 0.1 compared to wildtype expression at t = 0, with *q*-values representing Wald test *p*-values adjusted for false discovery rate. Legend abbreviations: BAHD, BAHD (BEAT, AHCT, HCBT, and DAT) acyltransferases; CESA, cellulose synthase; MLG, mixed-linkage glucans.

### SWIZ protein binding in the gene body is associated with dynamic changes in gene expression

Next, we investigated the direct binding targets of SWIZ protein by performing DNA affinity purification sequencing (DAP-seq) with whole genomic DNA. SWIZ interacted with 2,455 distinct positions in the genome (**Supplemental Table S15**). Those regions were significantly enriched for two E-box motif variants, (A/C)CAGNCTG(T/G) and (A/C)CAGCTG(T/G) (**Fig 4A**). Numerous binding sites were between the translational start and stop sites, with 21% of peaks in introns and 21% in exons (**Fig 4B**). Fewer than 10% of the peaks occurred in UTRs and the promoter (5 kb upstream of the 5 ‘UTR) and intergenic regions each accounted for 25%.

**Figure 4.**
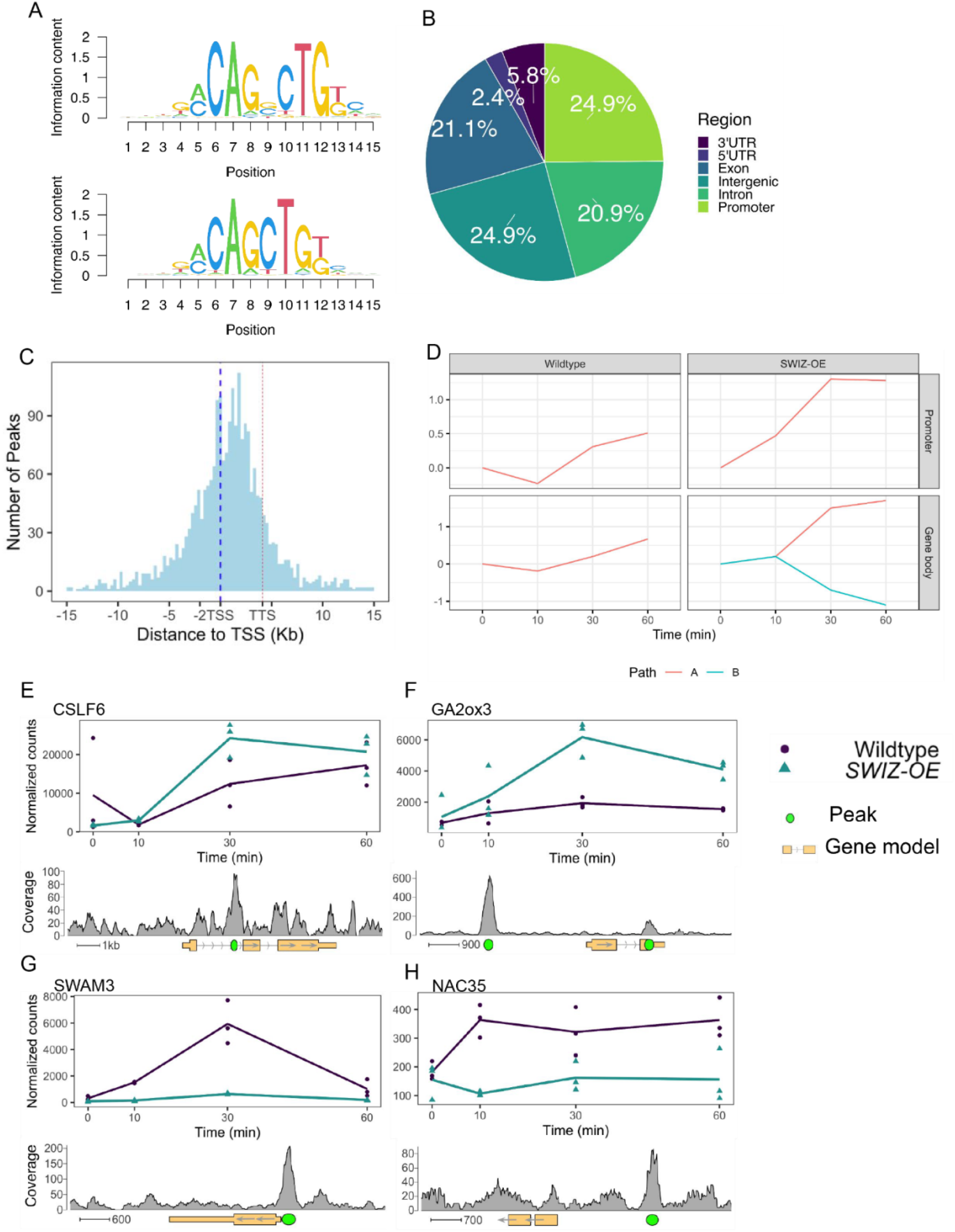
DNA affinity purification sequencing to determine SWIZ binding sites and differential expression in response to touch and *SWIZ-OE*. (A) Top two most statistically enriched sequence motifs in SWIZ binding sites. (B) Distribution of binding sites across genomic features, relative to primary transcripts of the *Brachypodium distachyon* annotation v 3.1. (C) Relative distribution of binding sites centered on the transcriptional start site (TSS, blue dashed line), transcriptional termination site (TTS, red dashed line) represents the average length of all annotated transcripts, approximately 4.5 kb away from the TSS. (D) Path determinations from iDREM time course analysis of differentially expressed genes that also have DAP-seq binding sites. Each line represents a set of genes with similar expression level patterns over the time course relative to time 0, pre-touch. Wildtype: promoter and gene body = 170 and 146 genes, respectively. *SWIZ-OE*: promoter = 163 genes, gene body A and B = 141 and 60, respectively. (E-H) Gene expression over time of selected genes with SWIZ binding sites. Line graphs are the average transcript abundance of three biological replicates for each time point. Binding site determined as peaks of sequence alignment. Scale bar unit is bases. Direction of transcription is shown with arrows on the gene model, 5’ and 3’ UTRs are depicted by narrowed rectangles on the gene model.

Thus, SWIZ protein is preferentially bound to an E-box like motif and often in the gene body (**Fig 4C**).

We next compared genes differentially expressed in the *SWIZ-OE* touch response RNA-seq time courses with the binding targets identified by DAP-seq (**Supplemental Table S5-9,15)**. Prior to touch, genes with promoter-SWIZ interactions were most often downregulated in *SWIZ-OE* relative to wildtype (**Table 1**). Following touch, SWIZ promoter binding targets were most often upregulated in *SWIZ-OE* in all three post-touch time points. Thus, SWIZ promoter binding was coincident with repression, but following a touch stimulus, activation was more prominent. This difference between untreated and touched was not as pronounced in gene body targets, with more upregulated genes 10 min following touch and more downregulated 30 and 60 min following touch. To further explore these trends, we conducted iDREM cluster analysis of the differentially expressed genes that were also SWIZ protein targets (**Fig 4D**). For both wildtype and *SWIZ-OE*, the network analysis revealed a pathway to increased gene expression by 60 min following touch. The network pathway trend for both promoter and gene binding targets in wildtype was immediate repression at 10 min followed by increased expression at 30 and 60 min post-stimulus. A unique pathway was observed among gene body binding targets in *SWIZ-OE*; transcript abundance was reduced following touch. Thus, SWIZ binding to a promoter or gene body was more strongly associated with increased expression except for gene body targets in touched *SWIZ-OE* plants, which were more strongly repressed.

**Table 1.** The number of SWIZ Dap-seq targets differentially expressed in wildtype plants after touch treatment relative to untouched wildtype The location of SWIZ DAP-seq binding, in either the promoter or gene body regions, is noted as is the direction of differential expression, up or down.

| Time after touch | Promoter |  | Gene body |  |
| --- | --- | --- | --- | --- |
|  | Up | Down | Up | Down |
| (min) | percent (count) |  |  |  |
| 0 | 42.0 (103) | 58.0 (142) | 48.0 (286) | 52.0 (310) |
| 10 | 57.0 (73) | 43.0 (55) | 53.0 (167) | 47.0 (148) |
| 30 | 59.4 (79) | 40.6 (54) | 41.2 (136) | 58.8 (194) |
| 60 | 64.2 (77) | 35.8 (43) | 45.7 (128) | 54.3 (152) |

Several vignettes stood out as examples of genes differentially expressed by touch, *SWIZ-OE* transgene, and being bound by SWIZ at various locations. The mixed glucan synthase *CSLF6* was bound in the first intron and upregulated by touch and *SWIZ-OE* (**Fig 4E**). *GA2ox3*, which inactivates bioactive GA, was bound by SWIZ protein in both the gene body and 3’UTR and was upregulated 30 min after touch in *SWIZ-OE* (**Fig 4F**). An example of gene body binding repression is *SWAM3* (Bradi1g30252), the closest ortholog to a wheat transcription factor induced by hypoxia, *TaMYB1* (Lee et al., 2006; Handakumbura et al., 2018) (**Fig 4G**). A membrane-associated transcription factor, NAC35, was bound by SWIZ in the promoter region and downregulated in SWIZ-OE (**Fig 4H**).

### *Cis*-regulatory sequences associated with mechanical stress, wounding, and cell wall synthesis are enriched among touch responsive genes

Touch-responsive genes in wildtype were analyzed for enrichment of putative *cis*-regulatory elements (CREs). We identified several sequences significantly enriched among touch-responsive transcripts (**Fig 5, Supplemental Fig S4, Supplemental Table S16-17**). Several of those have been previously described as touch responsive, including the Rapid Stress Response Element, the GCC boxes AGCCGCC and GCCGCC, a sequence referred to as both the E-box and G-box (CACGTG), CM2, AP2-like, GGNCCCAC site II element, P-box, and the GRF and FAR1 binding sites (Rushton et al., 2002; Walley et al., 2007; Doherty et al., 2009; Fernández-Calvo et al., 2011; Moore et al., 2022). Other putative CREs have not previously been identified as touch responsive. One CRE was exclusively enriched in touch-repressed genes at all time points, the TCP site II element TGGGC. A GATA-like binding site was enriched among 10 min repressed transcripts. The homeobox binding site motif was enriched among both induced and repressed genes. The CGCG-box was enriched among induced genes at all timepoints like RSRE, CM2, and FAR1. The two CREs associated with secondary cell wall thickening were also significantly enriched (Coomey et al., 2020); the VNS element among induced genes, and the AC element ACC(A/T)ACC with a unique profile. AC elements were enriched among repressed genes at 10 min, in both induced and repressed genes at 30 min, and among induced genes only at 60 min.

**Figure 5.**
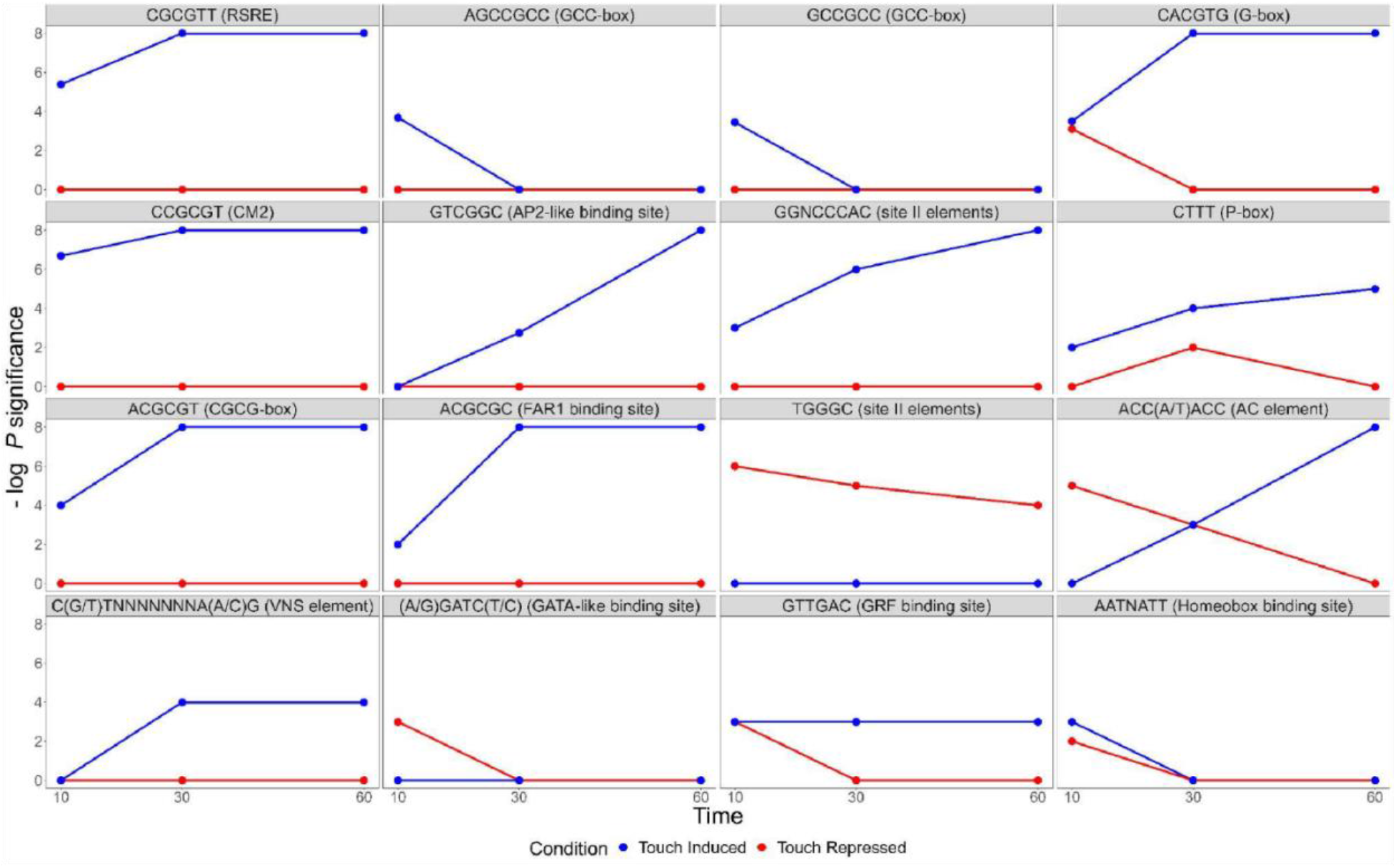
Sequence motifs enriched in the *cis*-regulatory regions of touch responsive *Brachypodium distachyon* genes. Negative log *p-*values for *cis*-elements, known and not known to be touch responsive. RSRE - Rapid Stress Response Element, FAR1 - FAR-RED impaired response1, GRF - Growth Regulating Factor, VNS - VND, NST/SND, SMB.

### Root morphology is altered by *SWIZ-OE* and mechanical treatment

Given the SWIZ protein mechanical response and regulatory influence, we then investigated how SWIZ might impact plant growth in response to touch. We challenged wildtype and *SWIZ-OE* roots with the same style of touch treatment used in the translocation and gene expression experiments, but repeated twice daily for a period of five days. In both touch and control conditions, *SWIZ-OE* roots were significantly shorter than wildtype, suggesting a dwarfing effect from *SWIZ* overabundance (**Supplemental Fig S5A-C**). Mechanical challenges to roots have been reported to impact root straightness, a trait that has also been described as being impacted in other bZIP studies (Van Leene et al., 2016; Zha et al., 2016). In control conditions, we observed that *SWIZ-OE* roots were significantly less straight than wildtype, while this was not observed in in response to touch (**Supplemental Fig S5D**). We further tested the mechanoresponse of *SWIZ-OE* roots by growing seedlings on plates with increasing degrees of plate angle at 10°, 20°, 30°, and 40° from vertical as a form of mechanostimulation (Oliva and Dunand, 2007; Zha et al., 2016) (**Supplemental Fig S6**). Wildtype *B. distachyon* roots displayed decreasing root length with increasing plate angle. *SWIZ-OE* roots were shorter than wildtype at all plate angles tested (**Supplemental Fig S6**). Root straightness did not show any significant differences.

### *B. distachyon* displays classic thigmomorphogenic phenotypes

We next investigated the effect of touch treatment on aboveground tissue. Wildtype plants were perturbed with a metal bar once every 90 min for either two or three weeks (**Supplemental Fig S7 and S8**). After the treatment period, all plants were allowed to recover and grow to senescence (**Supplemental Fig S8**). Two-week stressed plants were significantly shorter than control plants, and three-week stressed plants were shorter still (**Supplemental Fig S8A-B**).

Despite this difference in height, there was no significant difference in aboveground biomass (**Supplemental Fig S8C)**. Three-week stressed plants had significantly more branches, (**Supplemental Fig S8D**). Transverse stem cross sections in the third elongated internode and peduncle did not show a significant difference in cell wall thickness or phloroglucinol staining in response to touch (**Supplemental Fig S8E-G**).

### Stem height and cell wall thickening are affected by *SWIZ-OE* and touch treatment

To test the role of *SWIZ* in regulating thigmomorphogenesis and secondary cell wall development, we tested wildtype and *SWIZ-OE* under two weeks of mechanical perturbation as described above. In control conditions, there were no differences among genotypes in height, weight, or branching. Touch significantly shortened both wildtype and *SWIZ-OE* stems relative to control conditions, but not relative to each other (**Supplemental Fig S9-10**). Transverse sections of the stem were made in the peduncle, the last elongated stem internode where the touch treatment occurred during stem elongation, and stained with phloroglucinol-HCl. *SWIZ- OE* touched peduncles showed significantly thicker interfascicular fiber cell walls compared to untouched, but not significantly different from touched wildtype (**Supplemental Fig S10B-C**).

## DISCUSSION

Touch stimulus is generally an inhibitor of plant elongation growth, but promotes branching and radial expansion (Jaffe, 1973; Braam, 2004; Chehab et al., 2009). The majority of this work has been done in dicots, where increased radial growth has been associated with greater deposition of secondary wall forming cells, particularly in woody species such as poplar (Biro et al., 1980; Coutand et al., 2009; Börnke and Rocksch, 2018; Roignant et al., 2018; Niez et al., 2019). Our understanding of thigmomorphogenesis in grasses is limited, and mostly in the agricultural context of lodging (Shah et al., 2019). While these studies highlight the importance of stem strength and associated cell wall defects, they do relatively little to elucidate the mechanosensitive response. Recent work by Gladala-Kostarz et al (2020) describes grass touch response to both wind and direct mechanical treatment, with an emphasis on cell walls and stem anatomy. Touch treatment significantly increased lignin content and wall stiffness.

Specific gene expression patterns have become molecular hallmarks of plant touch response, most notably induction of the *TCH* and similar genes. Orthologs of wall modifying XTHs and CAM/CML signaling genes are upregulated in response to mechanical stimulation across species, and as we present here, in *B. distachyon*. Touch elicits major global changes in gene expression, with 2.5% to 10% of the genes assayed in *A. thaliana* and poplar differentially expressed (Lee et al., 2005; Pomiès et al., 2017; Van Moerkercke et al., 2019). In sorghum (*Sorghum bicolor*), leaf sheath imposes mechanical constraints on emerging buds, and removing this dramatically altered gene expression, with 42% of genes differentially expressed over a 9 h period (Liu and Finlayson, 2019). Recent work in cereal species, including oat, wheat, and barley, showed 2-5% differential gene expression within 2h of mechanostimulation of young leaves with a soft brush (Darwish et al., 2023). In our analysis we applied a *q*-value significance cutoff of 0.1, which accounts to some extent for the larger number of genes we find differentially expressed compared to other reports. An interactive application (https://hazenlab.shinyapps.io/swiztc) allows users to sort gene expression results by *q*-value, *p*- value, or log-2 fold change, and may be utilized by readers to make their own comparisons. Our results are similar to studies in other species in the strong induction of canonical TCH genes, along with biotic and abiotic stress related genes (Pomiès et al., 2017). Canonical touch responsive genes such as the orthologs of *TCH1-4* and related *XTH* and *CML* genes were upregulated immediately following stimulus. We also note the previously unreported touch- induction of GH17 family genes, as well as *CSLF6*, suggesting a role for MLG and β-1,3 glucan modification in grass touch response.

Touch-regulated expression of secondary cell wall related transcripts appears to differ between early and late timepoints. Upon touch, we observed immediate repression of many key cell wall biosynthetic enzymes, followed by significant upregulation one hour after stimulus. In poplar, no significant repression of these transcripts was reported, but terms related to wood or cell wall synthesis were also not enriched among touch-regulated transcripts until 24 or 72 hours after touch (Pomiès et al., 2017). In other cereals, genes related to cell wall polysaccharides were upregulated 25-60 min following touch while lignin biosynthetic genes down regulated (Darwish et al., 2023). While differences in mechanostimulation, tissue type, gene expression quantification, and temporal sampling schemes complicate direct comparisons of plant touch response experiments, together these data suggest that delayed induction of secondary cell wall transcripts is a common feature of plant touch responses.

### Thigmomorphogenesis and secondary cell walls

The timing of SWIZ nuclear accumulation following mechanostimulation was consistent with *A. thaliana* Group I bZIPs (Tsugama et al., 2014; Tsugama et al., 2016). However, by adopting a finer temporal sampling scheme, and quantifying nuclear signal sooner after touch (2 vs. 30 min), our results clarify the rapidity of this translocation. Furthermore, we show that the speed and magnitude of SWIZ nuclear accumulation is not diminished over successive touch treatments. Poplar transcriptional response to a second touch stimulus was significantly attenuated relative to a single treatment and trees given daily touch treatment over four days no longer showed touch-induced increases in their rates of radial growth (Martin et al., 2010; Pomiès et al., 2017). If *B. distachyon* is likewise desensitized to repeated touch, the repeatability of SWIZ translocation implies a mechanism downstream of touch perception and bZIP translocation in mediating that shift.

Although we did not observe a robust secondary wall response in touched stems, touch-responsive wall synthesis is still implicated by our transcriptomic data. Two CREs not previously associated with touch response, the AC and VNS elements, were identified in our touch-responsive and SWIZ-targeted datasets. These sites are bound by NAC and MYB proteins that regulate the thickening of secondary cell walls (Ohtani et al., 2011; Zhong et al., 2011; Kim et al., 2012; Zhong and Ye, 2012; Handakumbura et al., 2018; Olins et al., 2018; Tamura et al., 2019).

Prior to touch, almost the entire monolignol biosynthetic pathway was significantly downregulated in *SWIZ-OE* plants. Following touch, most of these genes were significantly upregulated. This activity is consistent with the compositional data presented by Gladala-Kostarz et al. (2020). This activation may be a result of direct binding of SWIZ, or an indirect effect from other transcriptional regulators like the NAC transcription factor *SWN5* that is significantly upregulated in *SWIZ-OE* following touch and capable of activating the full developmental program for secondary cell wall synthesis (Valdivia et al., 2013). The E-box (CANNTG) was initially described as a bHLH binding motif, many bZIP phylogenetic groups in *A. thaliana* have been shown to bind similar sequences (O’Malley et al., 2016). Using the same technique, we identified the SWIZ binding motifs as (A/C)CAGNCTG(T/G) and (A/C)CAGCTG(T/G), very similar to those bound by *A. thaliana* orthologs, although an ambiguous nucleotide in the core of one variant is not reported for other Group I bZIPs. Genes bound by SWIZ *in vitro* show both activation and repression following touch, suggesting a complex regulatory function for SWIZ. Furthermore, SWIZ direct binding sites were found in both promoter regions and gene bodies, and most often in genes activated by touch. SWIZ gene body binding targets tended to be repressed in *SWIZ-OE* plants following touch, without either being clearly associated with up or down regulation. AtVIP1 and AtbZIP29 have both been described as activators (Yin et al., 1997; Ringli and Keller, 1998; Pitzschke et al., 2009; Van Leene et al., 2016) while AtbZIP18 is described as a repressor (Tsugama et al., 2012; Gibalová et al., 2017). The directionality of SWIZ transcriptional control remains to be determined.

## CONCLUSIONS

Touch significantly remodeled the *B. distachyon* transcriptome, with notable changes in wall polysaccharide biosynthetic gene expression not previously reported and revealed an enrichment of secondary cell wall associated CREs, the AC and VNS elements. SWIZ mechanotropism shows similar dynamics to other bZIP proteins and the timing of this translocation is consistent with the enhanced touch-responsive gene expression in *SWIZ-OE*. Some of the data presented here relies on overexpression of *SWIZ*, which comes with certain limitations as to how directly the phenotypic and molecular results relate to SWIZ-specific function in wildtype plants. Further experimentation is needed to elucidate the bZIP regulatory network in response to touch and other stimuli.

## MATERIALS AND METHODS

### Phylogenetic analysis

Protein sequences described for *A. thaliana*, *B. distachyon*, and *O. sativa* (Liu and Chu, 2015) as Group II bZIPs were selected and searched against the most recent genome annotations in Phytozome v12.1 (https://phytozome.jgi.doe.gov). The *Nicotiana tabacum* homologs NtRSGa and NtRSGb were also added to the analysis. Protein sequences were aligned by MAFFT using the L-INS-I model (Katoh et al., 2019). A maximum-likelihood phylogenetic tree bootstrap resampling value of 1000 was generated in W-IQ-TREE (Trifinopoulos et al., 2016). All proteins in the phylogenetic analysis are described in **Supplemental File S1**.

### Plant transformation

Overexpression constructs and plant transformation carried out as previously described using accession Bd21-3 (Handakumbura et al., 2013). Bradi1g17700 coding sequence was amplified from cDNA and cloned into the pOL001 ubigate ori1 binary expression vector (Handakumbura et al., 2018) to make the *SWIZ-OE* trangene. The coding sequence without stop codon was amplified and cloned in frame with the enhanced GFP coding sequence in the pOL001 vector to generate *SWIZ:GFP-OE*. For both constructs, three independent events were analyzed with no phenotypic difference between them for height or translocation dynamics among the *SWIZ:GFP-OE lines*.

### Translocation assay

Bd21-3 seeds were surface sterilized and grown vertically on 1X MS media, pH 5.7, without sucrose for 6d at 28°C in the dark. After 6 d, seedlings were moved to treatment plates containing 1X MS media, pH 5.7.

All observations were made on a Nikon A1R scanning confocal microscope using a Plan Apo 10x 0.5NA objective and PMT detector. Root areas were located by eye using transmitted light and then imaged with excitation at 488 nm and emission captured at 510-530 nm. Roots were imaged for 30 min pre-treatment, with images captured every 2 min.

To elicit the touch response, the observed root region was gently probed 5 times in ∼5 sec with a blunt probe while observing through the eyepiece (**Supplemental Fig S7**). Images were captured for 60-90 min post treatment. For experiments with multiple stimulus events, the timelapse sequence was paused and roots were probed as described for the relevant stimulus events.

Analysis of GFP signal was done using the Nikon NIS Elements Advanced Research V5 software package. Nuclear regions were thresholded for intensity and particle size to identify regions of interest. Fluorescence intensity was calculated for each ROI and averaged at each timepoint.

### Thigmomatic construction and operation

The Thigmomatic is a basic robotic device that sweeps plants with a metal bar at regular intervals to elicit a touch response. The device was constructed from aluminum V-Slot linear rail (Openbuilders Partstore, Monreoville, NJ) and bracket joints for the upright supports (20x20 mm), cross bars (20x20 mm), and tracks (20x40 mm). Two gantry carts ride along the 20x40 mm V-Slot linear rails, connected by a 6.35 mm diameter metal rod bolted to the carts. Their movement is powered by a belt driven linear actuator system using a NEMA 17 stepper motor with a 12V 18W AC/DC power supply. Motor function is controlled by a Raspberry Pi 3B microcomputer with stepper motor HAT (Adafruit Industries, New York). The Thigmomatic was programmed to cover a specified distance in one direction once every 90 min.

### Transverse stem sections and histology

Internode segments of the main stem were embedded in 8% agarose. A Leica VT1000 Vibratome was used to make 55 µm thick transverse sections. Histochemical staining with phloroglucinol-HCl was done as previously described (Matos et al., 2013). Images were obtained at 4, 10, and 20X using a Nikon Eclipse E200MV R light microscope and PixeLINK 3 MP camera. Cell wall thickness was quantified for interfascicular fiber cells separated by one cell layer from the mestome cells on the phloem side of major vascular bundles. Using ImageJ, lines were drawn across two adjoining walls divided by two yielding one cell wall width.

### RNA extraction and quantification and analysis

Seedlings were grown for 6 d on vertical 1X MS agar plates. Touch treatment was performed as described above using a metal probe along the entire length of the root. Untouched samples were collected immediately before touch treatment and touched samples collected 10, 30, and 60 min post-treatment. Three roots were pooled per biological replicate, and RNA extracted by Qiagen RNeasy Plant Mini Kit with on-column DNA digestion with RNase-free DNase I (Qiagen).

Strand-specific libraries were prepared using the Illumina TruSeq kit. Libraries were sequenced using Illumina technology and processed as previously described (MacKinnon et al., 2020).

Briefly, quality was checked with FastQC (Andrews, 2010), aligned to the Bd21 reference genome (v3.1) using HiSat2 (Kim et al., 2015), then assembled and quantified using StringTie (Pertea et al., 2015). Transcripts were normalized and differential expression tested using the likelihood ratio test from the R (v3.6.0) package DESeq2 (Love et al., 2014). Benjamini-Hochberg *p*-value adjustments were applied to account for multiple testing with a significance cutoff of 0.1 and 30,380 of 34,310 reference genes had non-zero read counts after normalization, with an average mapping percentage of 97.8% for all libraries, as determined by SAMtools (Li et al., 2009). Specific treatment contrasts (i.e., wildtype vs *SWIZ-OE*) were compared by Wald test from DESeq2 (Love et al., 2014). Statistical enrichment of gene families was assessed using Fisher’s exact test. Raw read data was deposited in the European Nucleotide Archive for public access (Accession no.: E-MTAB-10084).

### DNA affinity purification sequencing

DNA affinity purification was carried out as previously described (Handakumbura et al., 2018). In brief, transcription factor coding sequences were HALO tagged and mixed with Bd21 genomic DNA for *in vitro* binding. Protein-DNA was crosslinked, fragmented, immunoprecipitated using the HALO antibody, barcoded, and sequenced. Reads were mapped to the Bd21 genome using HiSat2 (Kim et al., 2015) to identify binding target loci. Peak calling and motif analysis was done using HOMER v4.10 (Hypergeometric Optimization of Motif EnRichment) suite (Heinz et al., 2010). Motif enrichment was calculated against the hypergeometric distribution; the significance threshold was set to *p* < 0.05. The nearest annotated gene to a bound peak was used for GO analysis. Raw read data were deposited in the European Nucleotide Archive for public access (Accession no.: E-MTAB-10066).

### Gene Ontology analysis

Phytozome was used to find orthologs for all *B. distachyon* v3.1 genes as the reciprocal best match to *A. thaliana* TAIRv10 protein sequences. Arabidopsis gene identifiers were submitted to g:Profiler (Raudvere et al., 2019) for KEGG and Wiki pathway enrichment analysis.

### iDREM network analysis

Probabilistic graphical models that predict diverging gene expression paths were generated using iDREM (Ding et al., 2018). Briefly, this software applies an input-output hidden Markov model to time course gene expression data overlaid with static regulatory information, in this case SWIZ DAP-seq protein-DNA interactions. GO analysis, described above, was performed for each path identified by iDREM.

### *Cis*-regulatory sequence analysis

Differentially expressed genes following touch were categorized as increasing or decreasing in transcript abundance at each time point. Homer v.4.10 identified regulatory sequences in the 1000 bp upstream of the transcriptional start site of differentially expressed genes previously identified in *A. thaliana* DAP-seq analysis (Heinz et al., 2010; O’Malley et al., 2016). We also applied the growing k-mer approach to identify CREs (Moore et al., 2022). In brief, touch-responsive genes were divided into six groups: up or down regulated each timepoint. The 1000 bp upstream of differential expressed genes were searched for all possible 6-mers, plus one additional base (A,T,C,G), and then tested for *p*-value shift. If the *p*-value was lower, the 7- mer(s) was kept and grown further up to 12-mer length. The Tomtom tool in MEME Suite 5.4.1 found similarities between significant CREs and the *A. thaliana* DAP-seq database, with a false discovery rate cutoff of < 0.01.

### Root touch experiment

Bd21-3 seeds were surface sterilized and plated on 1x MS, pH 5.7, containing 0.05 % MES as a buffering agent and 1% plant agar (Gold Bio). Seeds were stratified on plates in the dark at 4 ℃ for 2 days and then transferred to a Percival PGC-15 growth chamber with day/night conditions of 16h light at 24 ℃ and 8h dark at 18 ℃, respectively and grown at a ∼10° angle from vertical. After 2 days, a touch location was designated by selecting the lower of 1 cm up from the root tip or 1 cm down from the seed and marked on the plate. For 5 days, this marked spot was treated twice daily, two hours before and two hours after chamber midday (ZT6 and ZT10). Touch treatment consisted of five firm presses with the side of a sterile pipette tip.

### Root length and straightness measurement

Plates were photographed at the conclusion of the experiment. Semi-automated measurements of root length were performed using the Smart Roots plugin for ImageJ (Lobet et al., 2011).

Straightness was quantified as described in (Swanson et al., 2015); the straight line distance between root tip and the base of the seed was measured, and this value was divided by the traced length of the root.

## SUPPLEMENTAL MATERIALS

Supplemental Table S1. Cluster 56 from network analysis of gene expression atlas (Sibout et al. 2017) includes Bradi1g17700.

Supplemental Table S2. Differentially expressed genes in wildtype root tissue 10 min after touch treatment.

Supplemental Table S3. Differentially expressed genes in wildtype root tissue 30 min after touch treatment.

Supplemental Table S4. Differentially expressed genes in wildtype root tissue 60 min after touch treatment.

Supplemental Table S5. Differentially expressed genes in SWIZ-OE root tissue prior to touch treatment (0 min).

Supplemental Table S6. Differentially expressed genes in SWIZ-OE root tissue 10 min after touch treatment.

Supplemental Table S7. Differentially expressed genes in SWIZ-OE root tissue 30 min after touch treatment.

Supplemental Table S8. Differentially expressed genes in SWIZ-OE root tissue 60 min after touch treatment.

Supplemental Table S9. Overlap of differentially expressed genes in wildtype and SWIZ- OE root tissue before touch (0 min) and 10, 30, and 60 min after touch treatment.

Supplemental Table S10. Count of differentially expressed genes in wildtype and SWIZ-OE root tissue 10, 30, and 60 min after touch treatment and the count of differentially expressed genes shared between genotypes at each timepoint.

Supplemental Table S11. Calmodulin and calmodulin-like touch responsive gene expression in wildtype Bd21-3 root tissue.

Supplemental Table S12. XTH and XTH-like touch responsive gene expression in wildtype root tissue.

Supplemental Table S13. Glycoside hydrolase touch responsive gene expression in wildtype root tissue.

Supplemental Table S14. GO analysis of genes in iDREM identified pathways derived from differential gene expression following touch in wildtype and SWIZ-OE root tissue.

Supplemental Table S15. DAP-seq results of SWIZ genome binding locations.

Supplemental Table S16. All putative CREs enriched in differentially expressed genes at each timepoint identified using the k-mer approach.

Supplemental Table S17. CREs described for Arabidopsis thaliana in O’Malley et al. 2016 that match CREs identified using the k-mer approach.

---

**Supplemental Figure S1.**
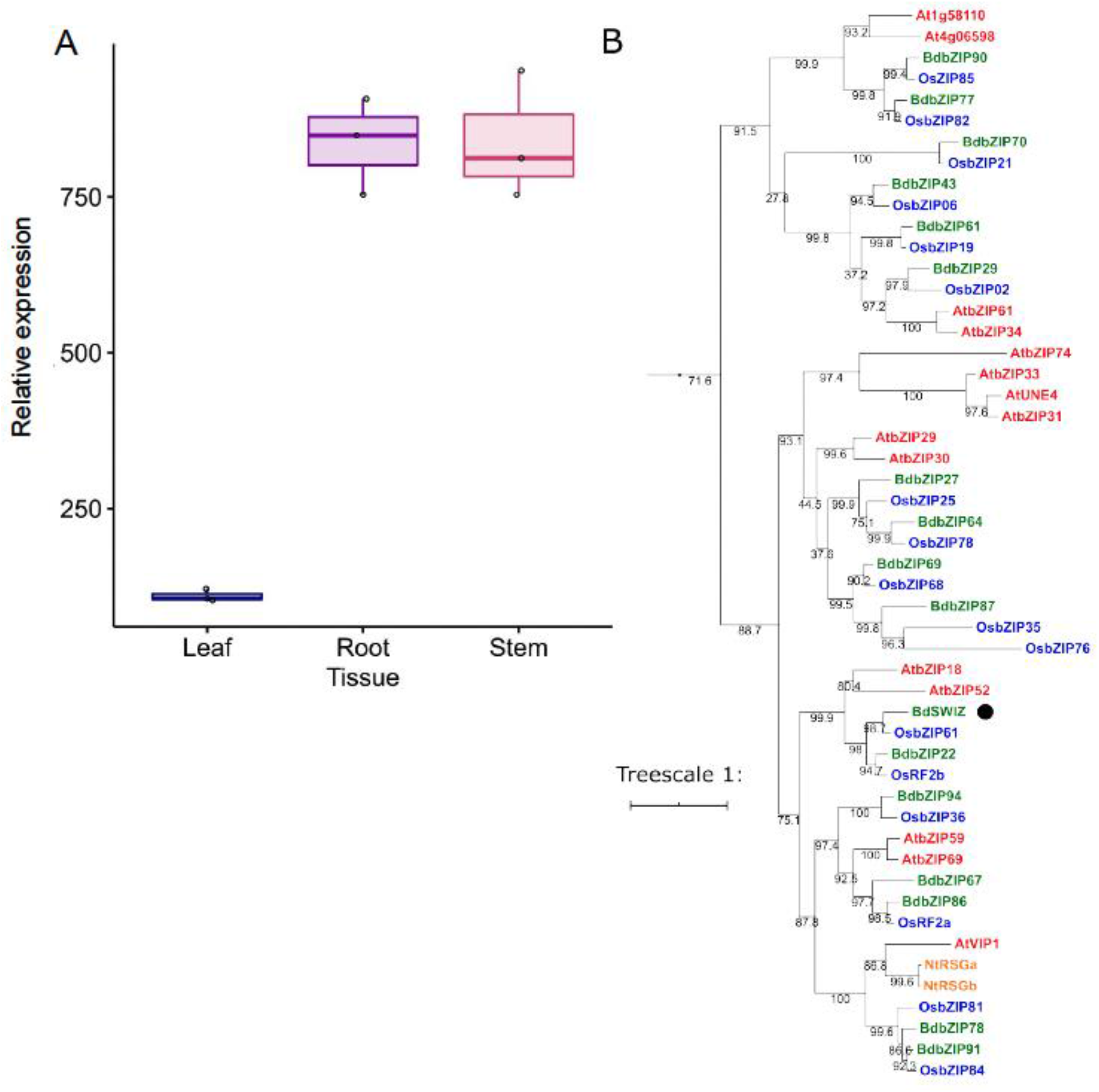
SWIZ is a Group I bZIP highly expressed in maturing stem and root. (A) *SWIZ* transcript abundance in *Brachypodium distachyon* leaf, root, and stem tissue measured by microarray. Mean +/- standard deviation of three biological replicates. (B) Phylogeny analysis of amino acid sequences from *B. distachyon* (green), *Oryza sativa* (blue), *Arabidopsis thaliana* (red), *Nicotiana tabacum* (orange) shows *SWIZ* (black dot). The phylogeny was reconstructed using maximum likelihood analysis with a bootstrap resampling value of 1000. The numbers labeled on the branches are the posterior probability to support each clade. The protein sequences, full locus IDs, and plant species with genome source are provided in Supplemental File S1.

**Supplemental Figure S2.**
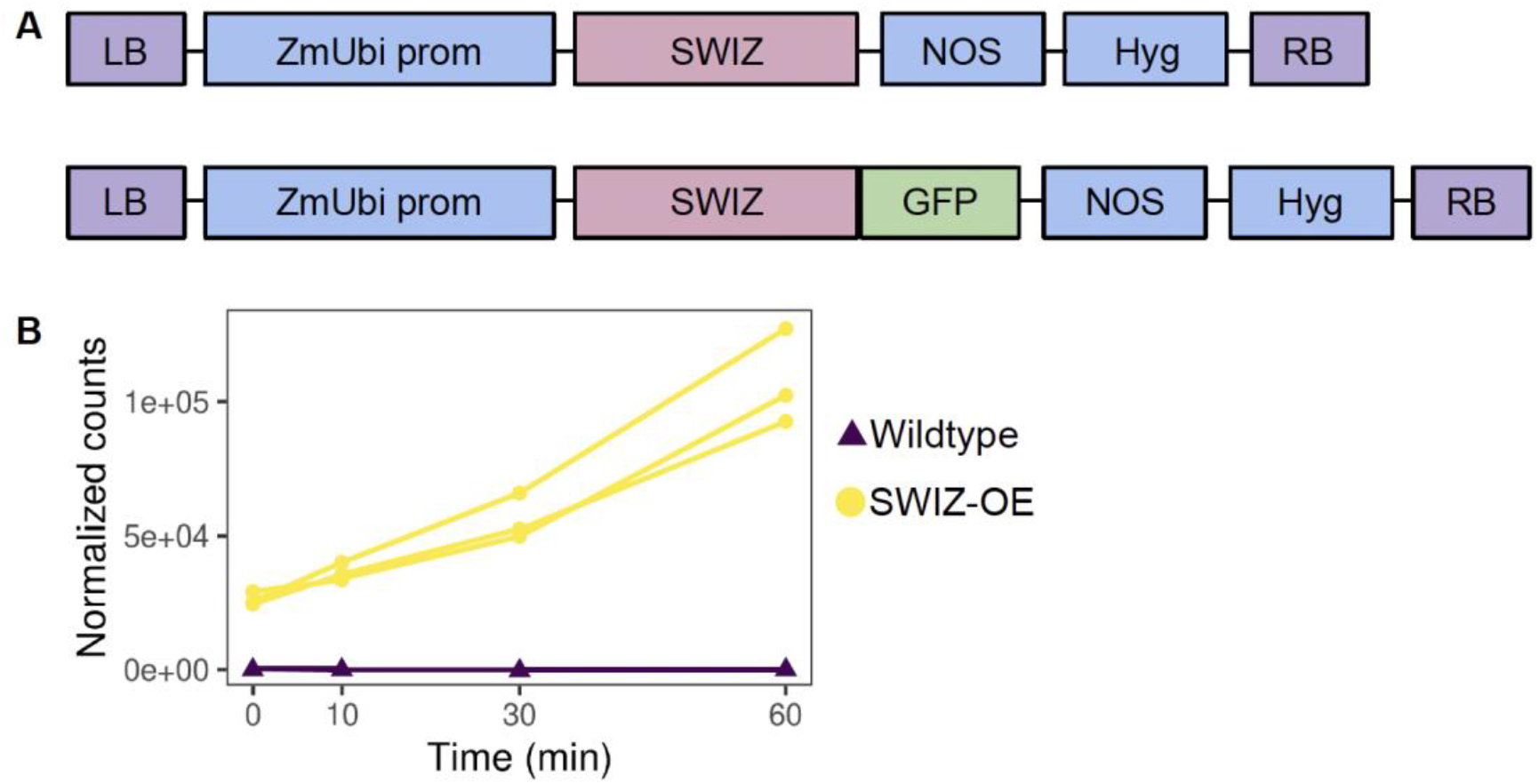
Description of *SWIZ* transgenic reagents. (A) Diagram of *SWIZ* overexpression transgenes. The *ZmUbi* promoter was used to drive expression of the *SWIZ* coding sequence, either alone or fused in frame with *eGFP*. (B) Transcript abundance of *SWIZ* measured by RNA-sequencing. Root tissue was sampled just prior to touch (t = 0), and at 10, 30, and 60 min following touch treatment in wildtype and *SWIZ-OE.* LB, left border; *ZmUbi* prom, maize ubiquitin promoter; Hyg, hygromycin phosphotransferase gene; NOS, nopaline synthase terminator; RB, right border.

**Supplemental Figure S3.**
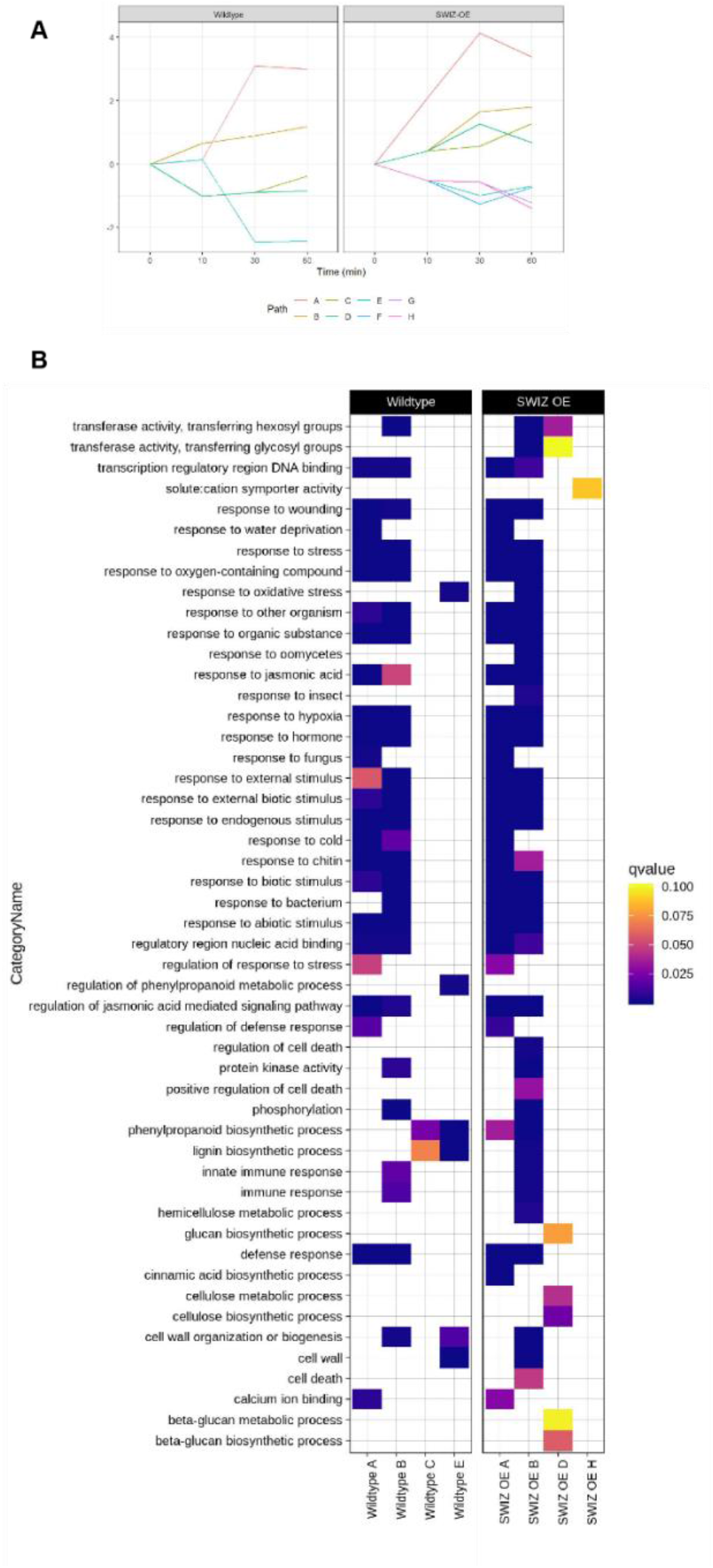
Network analysis of differential gene expression paths and enriched GO terms. (A) Path determinations from iDREM time course analysis. Each line represents a set of genes with similar expression level patterns over the time course relative to time 0, pre-touch. (B) Heatmap of selected GO term enrichment of genes from iDREM path analysis. Color indicates *q*-value score, with a cutoff of 0.1. Terms without a colored block are not significantly enriched in that path.

**Supplemental Figure S4.**
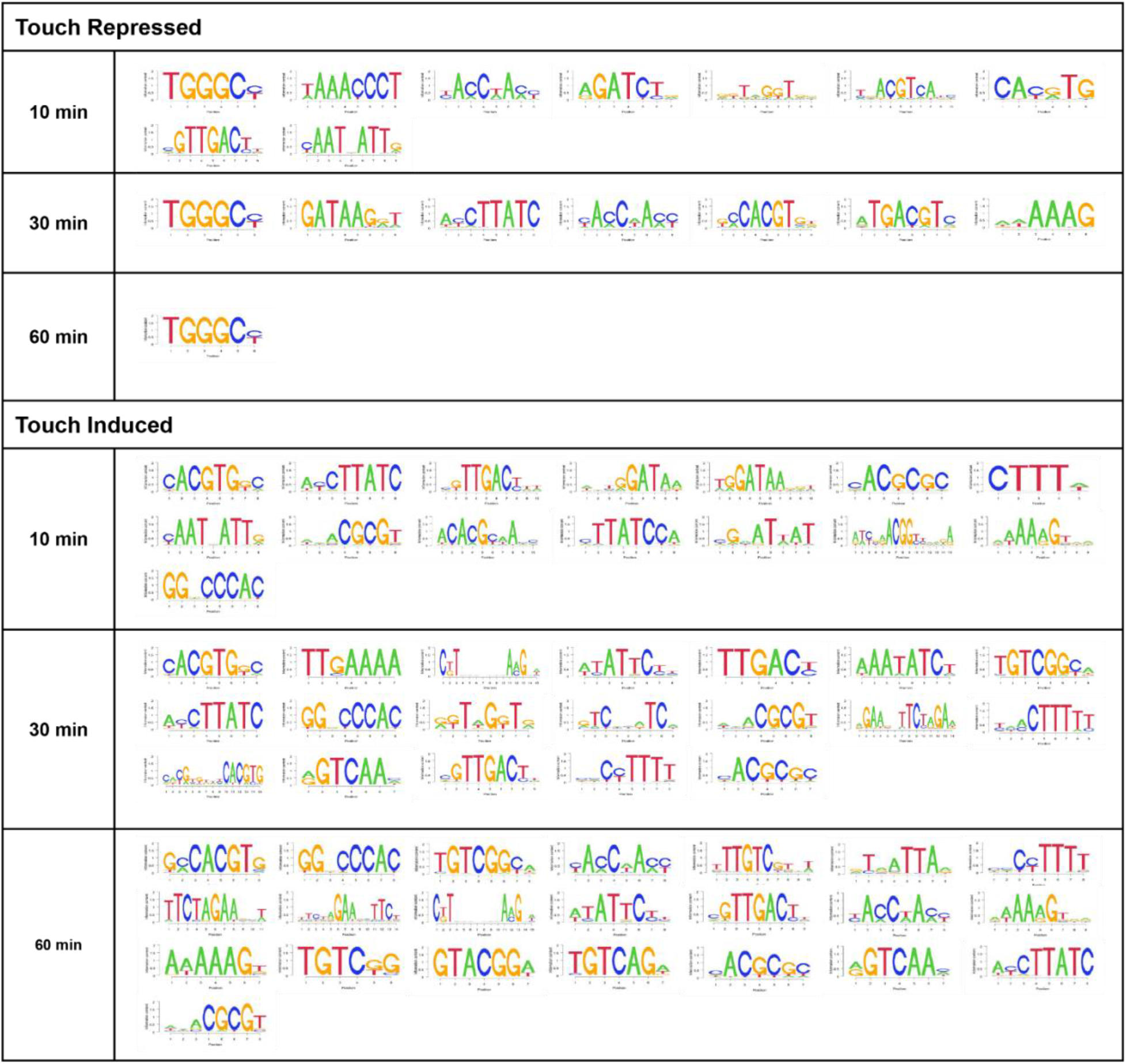
Brachypodium distachyon cis-regulatory sequences most overrepresented at each time course after touching. Each nucleotide sequence is a position probability matrix motif derived from DNA-affinity purification sequencing and identified as enriched under different time courses. The height of the letter at each position is proportional to the probability of a given nucleotide.

**Supplemental Figure S5.**
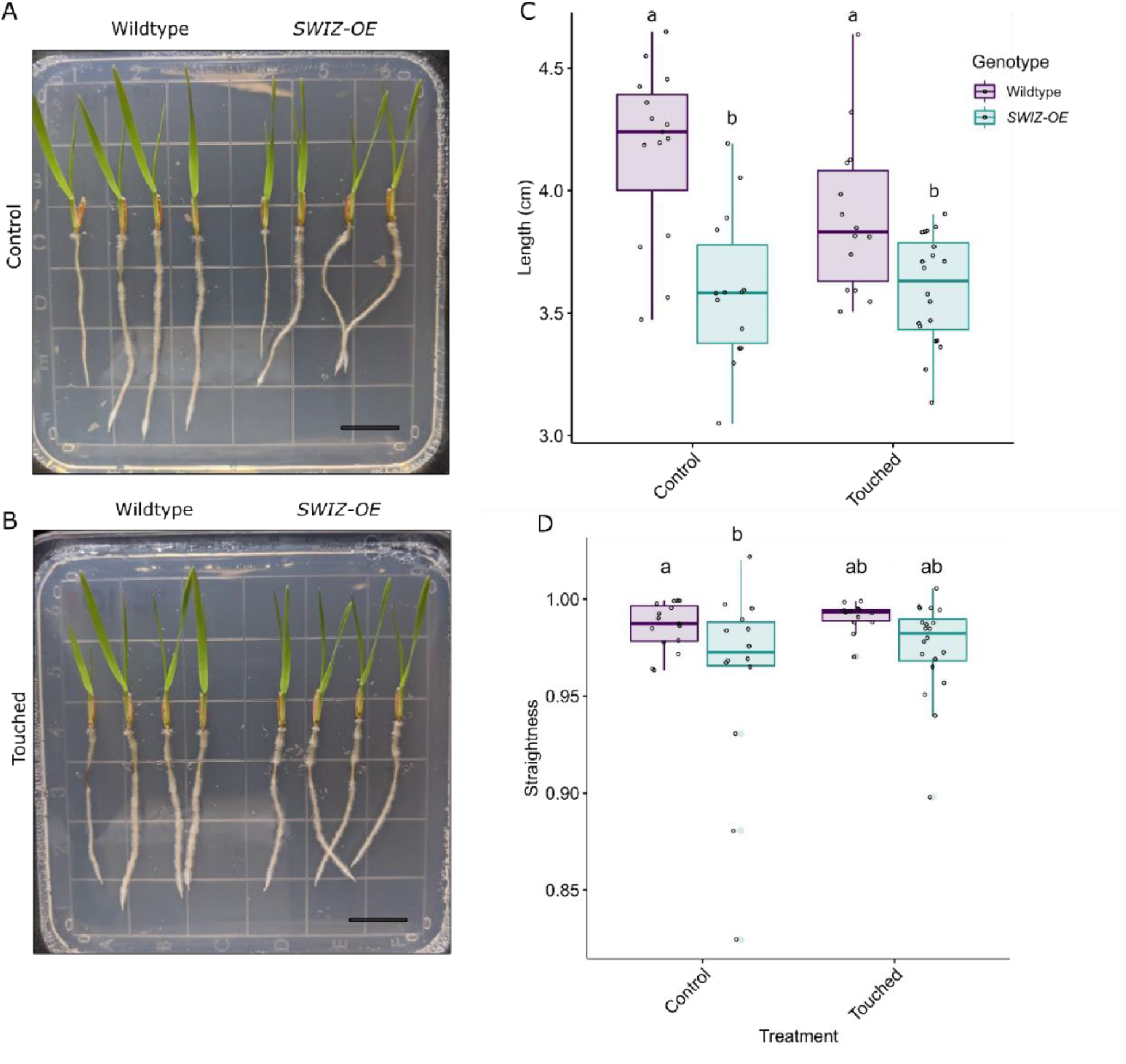
***SWIZ-OE* roots are shorter than wildtype with no significant difference in straightness in response to direct touch.** Wildtype and *SWIZ-OE* seedlings grown on plates under control (A) or touched (B) conditions for five days. Touched plants were probed with a pipette tip twice a day. (C) Quantification of root length in control and touched conditions. Significance denoted by compact letter display reflecting Tukey HSD adjusted *p-* values < 0.05. (D) Quantification of root straightness in control and touch conditions. Significance denoted by * reflecting Wilcoxon sign-ranked test for non-parametric data with *p*- value < 0.05, ns = not significant. Scale bar = 1 cm.

**Supplemental Figure S6.**
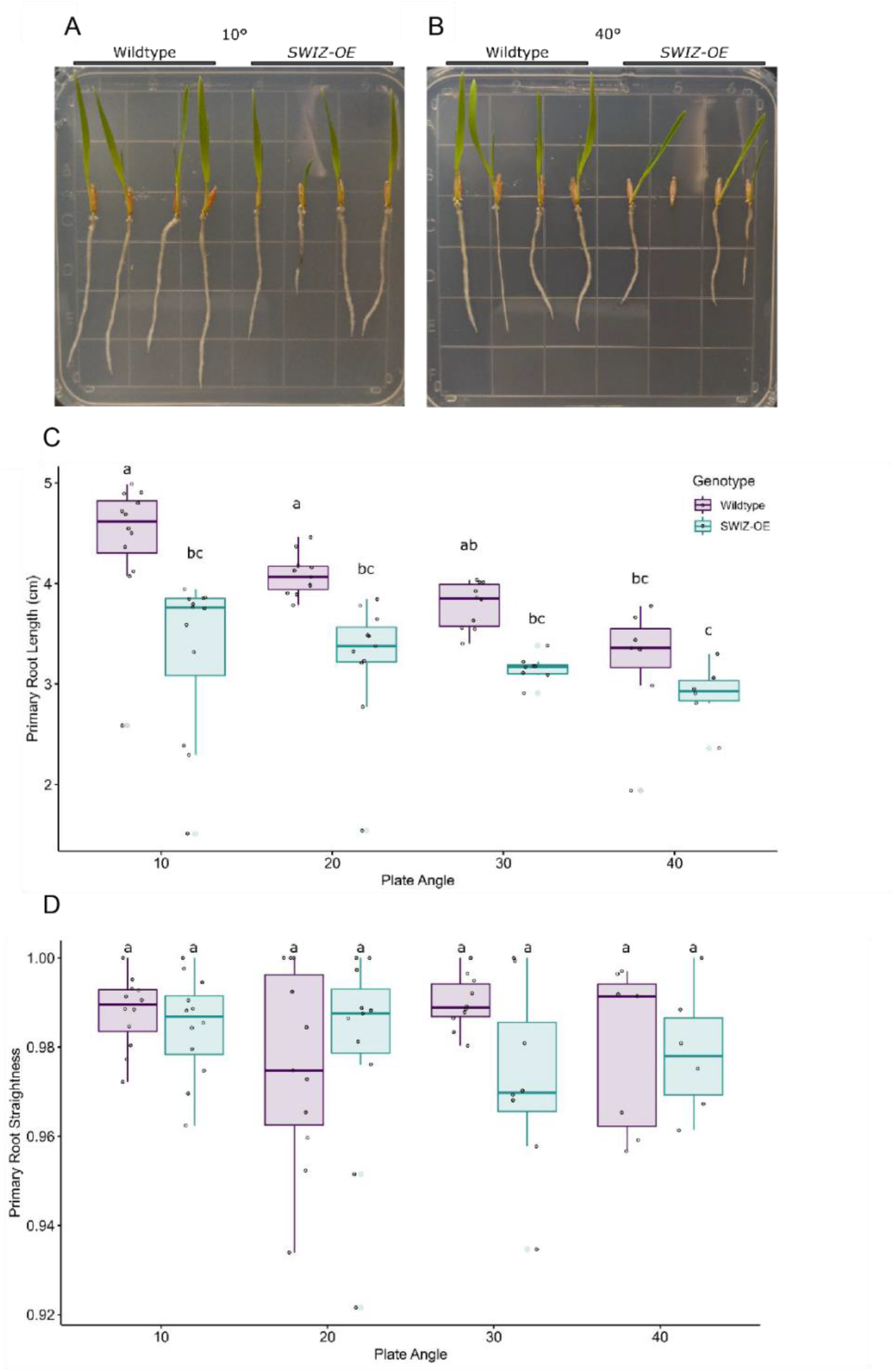
Wildtype roots shorten with increasing plate angle, while *SWIZ- OE* roots are consistently short, with no significant difference in straightness. Wildtype and *SWIZ-OE* seedlings grown on plates at (A) 10°, 20°, 30°, and (B) 40° incline from vertical. (C) Quantification of root length. (D) Quantification of primary root straightness. Significance denoted by compact letter display reflecting Tukey HSD adjusted *p-*values < 0.05. Scale bar = 1 cm.

**Supplemental Figure S7.**
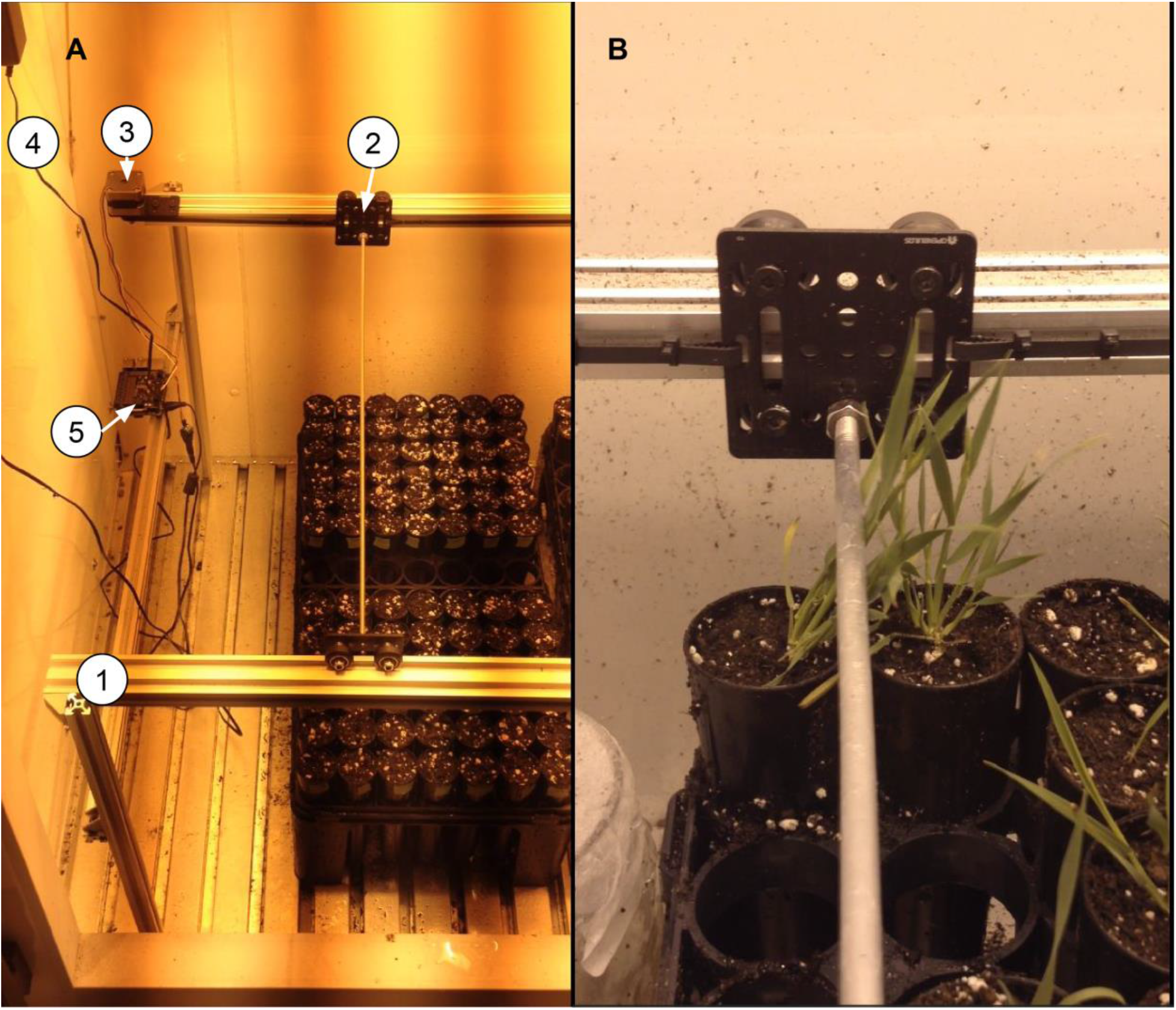
Thigmomatic, an automated system to administer touch stimulus. (A) Overview of Thigmomatic inside a Percival PGC-15 growth chamber showing linear rail based frame (1), gantry carts (2), NEMA17 stepper motor (3), 12V 18W AC/DC power supply (4), and Raspberry Pi 3b microcomputer (5). (B) Thigmomatic making contact with a *Brachypodium distachyon* plant.

**Supplemental Figure S8.**
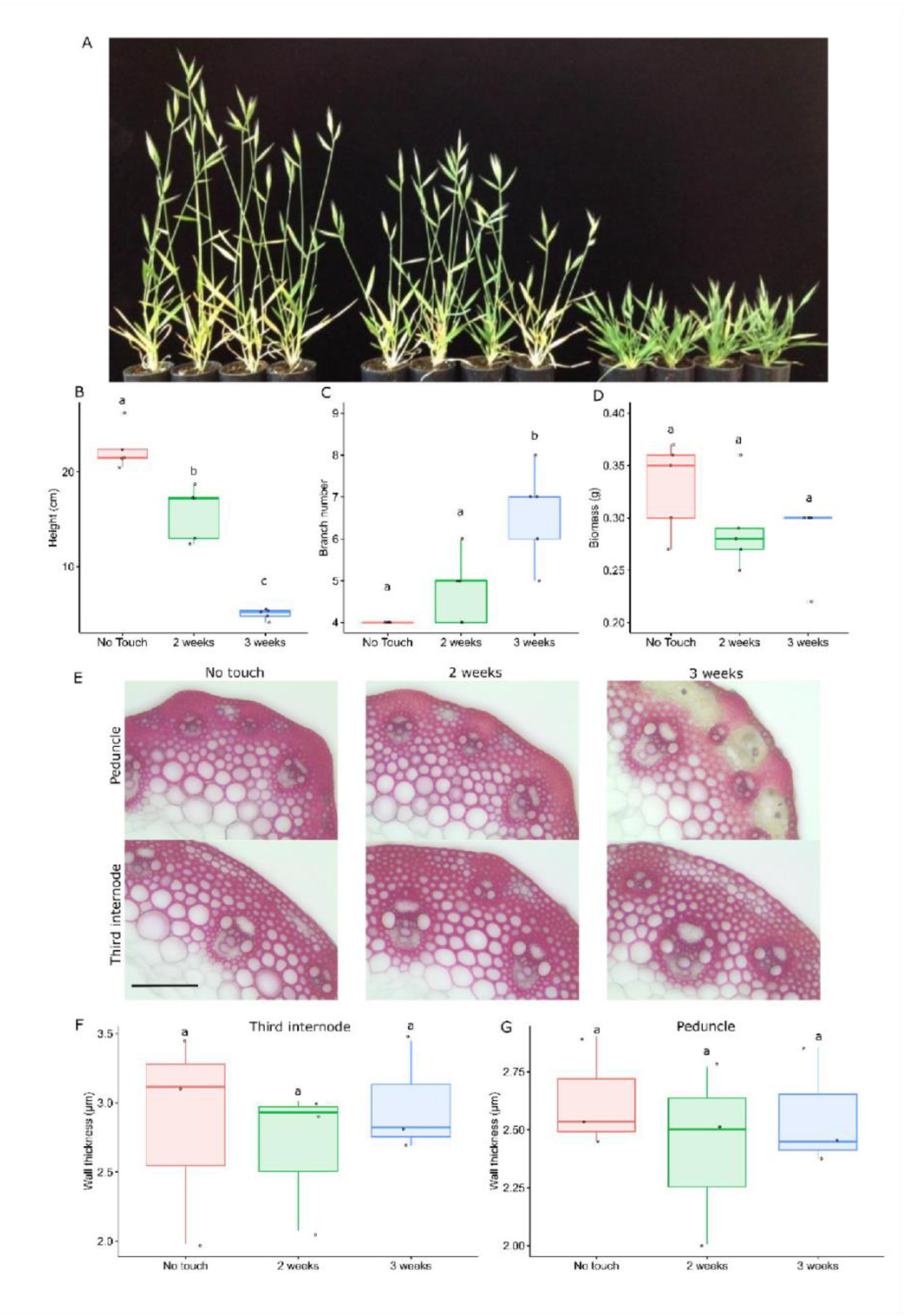
Brachypodium distachyon displays classic thigmomorphogenic phenotypes. One-week-old wildtype plants were treated with touch stimulus every 90 min. (A) Left to right, plants that experienced no stress, two weeks stress, three weeks stress and were then allowed to recover, imaged one week after the end of treatment (B) height, (C) aboveground non-grain biomass weight, and (D) branch number were measured at senescence. n = 4-5 plants per treatment. Significance denoted by compact letter display reflecting Tukey HSD adjusted *p*- values < 0.05. (E) Transverse sections of the peduncle or third internode were taken from control, 2 week stressed, and 3 week stressed plants and stained with phloroglucinol-HCl. (F) Quantification of interfascicular fiber wall thickness. Scale bar = 100 µm. n = 3 plants per treatment.

**Supplemental Figure S9.**
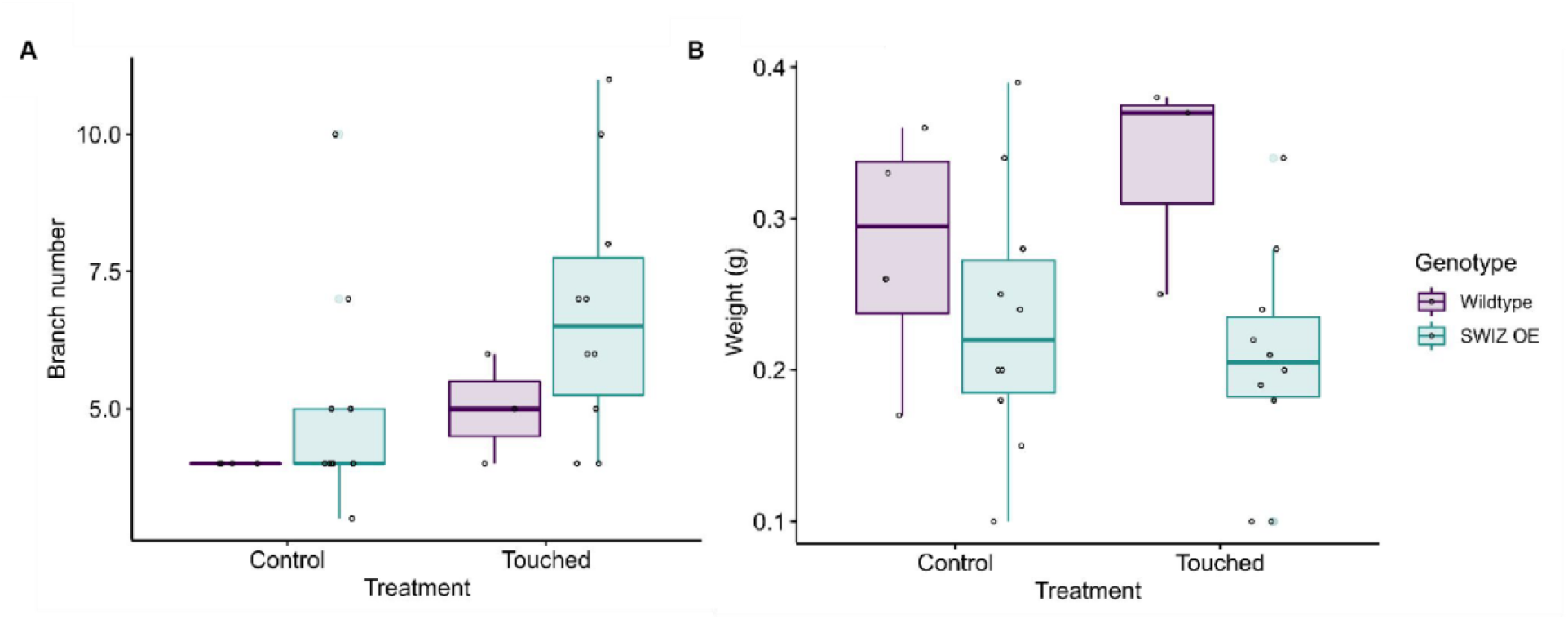
Biomass and branching in SWIZ-OE under touch treatment shows no difference from wildtype. Wildtype and SWIZ-OE plants were placed under control conditions or received two weeks of mechanical stress every 90 min. After senescence, branch number (A) and aboveground biomass weight (B) were quantified. Significant differences were not observed following ANOVA and Tukey HSD testing.

**Supplemental Figure S10.**
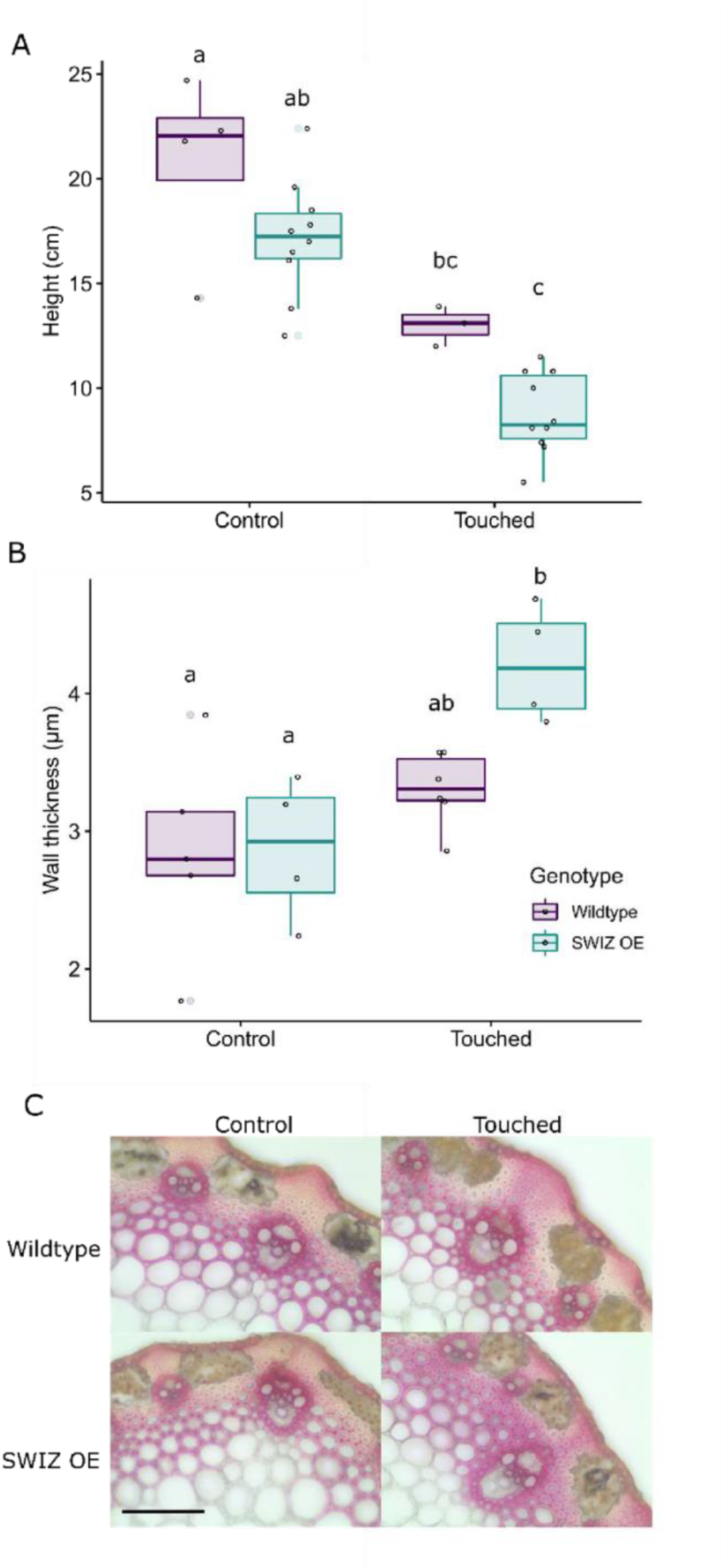
*SWIZ-OE* marginally enhanced thigmomorphogenic response in stems. Wildtype and *SWIZ-OE* plants were placed under control conditions or received two weeks of mechanical stress every 90 min. (A) Quantification of main stem height at senescence. (B) Quantification of interfascicular fiber wall thickness in transverse cross sections of the peduncle. n = 4 to 6 plants per genotype, per treatment. (C) Representative cross sections stained with phloroglucinol-HCl. Scale bar = 100 µm. Significance denoted by compact letter display reflecting Tukey HSD adjuste*d p-*values < 0.05.

**Supplemental Figure S11.**
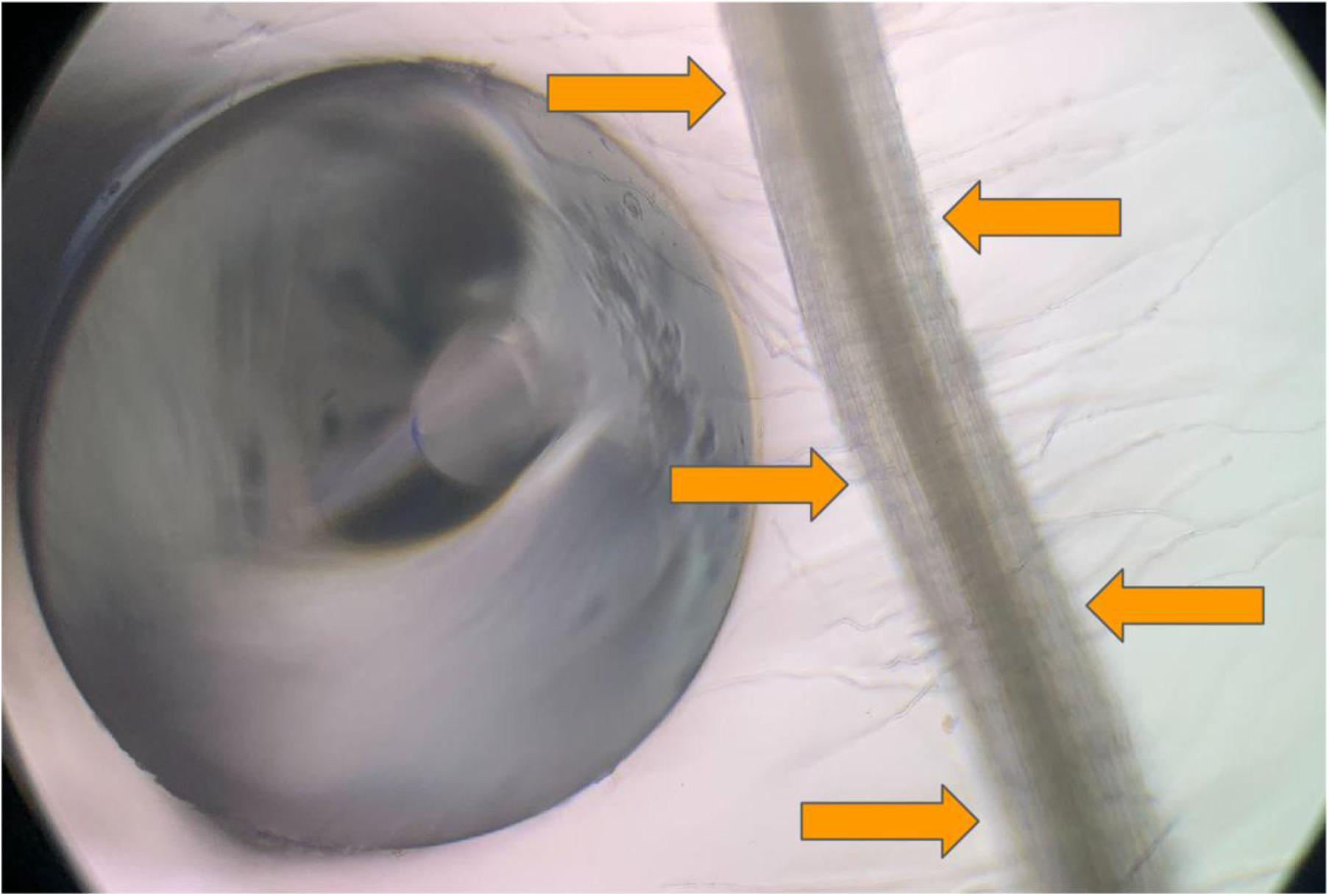
Method of root touch treatment. A blunt probe formed from a glass Pasteur pipette was used to gently tap on the root five times in areas like those indicated by the orange arrows.

Supplemental File S1. Gene names and amino acid sequences used in the SWIZ bZIP Group I phylogenetic analysis.

Supplemental File S2. Movie of *GFP-OE* root images captured for 60-90 min post touch treatment.

Supplemental File S3. Movie file of a *SWIZ*:*GFP-OE* root images captured for 60-90 min post touch treatment.

## ACCESSION NUMBERS

AtbZIP18 (At2g40620), AtbZIP29 (At4g38900), AtbZIP52 (At1g06850), AtTCH1/AtCaM2 (At2g41110), AtTCH2/AtCML24 (At5g37770), AtTCH3/CML12 (At2g41100), AtTCH4/AtXTH22 (At5g57560), CAD1 (Bradi3g17920), CESA4 (Bradi4g28350), COMT6 (Bradi3g16530), CSLF6 (Bradi3g16307), GA2ox3 (Bradi2g50280), SWAM3/MYB44 (Bradi1g30252), NAC35 (Bradi2g28020), NtRSGa (Niben101Scf01150g00005), NtRSGb (Niben101Scf02191g00014), SWIZ (Bradi1g17700), VIP1 (At1g43700).

## DATA AVAILABILITY STATEMENT

The data that support the findings of this study are openly available at E-MTAB-10066 and E-MTAB-10084 in the European Nucleotide Archive and https://hazenlab.shinyapps.io/swiztc.

